# Single-Cell Amplicon Sequencing Reveals Community Structures and Transmission Trends of Protist-Associated Bacteria in a Termite Host

**DOI:** 10.1101/299255

**Authors:** Michael E. Stephens, Daniel J. Gage

## Abstract

The hindgut protists of wood-feeding termites are usually colonized by prokaryotic symbionts. Many of the hurdles that have prevented a better understanding of these symbionts arise from variation among protist and termite host species and the inability to maintain prominent community members in culture. These issues have made it difficult to study the fidelity, acquisition, and differences in colonization of protists by bacterial symbionts. In this study, we use high throughput amplicon sequencing of the V4 region of 16S rRNA genes to determine the composition of bacterial communities associated with single protist cells of six protist species, from the genera *Pyrsonympha*, *Dinenympha*, and *Trichonympha* that are present in the hindgut of the termite *Reticulitermes flavipes.* By analyzing amplicon sequence variants (ASVs), the diversity and distribution of protist-associated bacteria was compared within and across these six different protist species. ASV analysis showed that, in general, each protist genus associated with a distinct community of bacterial symbionts which were conserved across different termite colonies. However, some ASVs corresponding to ectosymbionts (Spirochaetes) were shared between different *Dinenympha* species and to a lesser extent with *Pyrsonympha* and *Trichonympha* hosts. This suggested that certain bacterial symbionts may be cosmopolitan to some degree and perhaps acquired by horizontal transmission. Using a fluorescence-based cell assay, we could observe the horizontal acquisition of surface-bound bacteria. This acquisition was shown to be time-dependent, involve active processes, and was non-random with respect to binding locations on some protists.

## Introduction

The lower termite *R. flavipes* harbors symbionts from the three domains of life, all of which make significant contributions to the digestion of lignocellulose and provisioning of essential nutrients. These symbionts include uncultivated hindgut protists of two eukaryotic taxa, Oxymonadida (order) and Parabasalia (phylum or class) (1,2). Most, and perhaps all, of these protists are colonized by both endo- and ectosymbionts belonging to various bacterial taxa (2). These protist-associated bacteria often exhibit complex community structures and occupy different sites on and within their unicellular hosts (2).

Previous studies have shown that Oxymonadida protists in *Reticulitermes speratus* are co-colonized with *Treponema* ectosymbionts from two distinct phylogenetic clusters (Termite *Treponema* clusters I and II) (3,4) as well as a member of the Bacteroidales (5). These three ectosymbiotic lineages attach by one cell pole to the plasma membrane of their host (6) and exhibit intermixed colonization (5). Other known ectosymbionts include a *Desulfovibrio* species which embeds in the membrane of its host, *Trichonympha* (7). Functional and genomic data regarding the nature of these symbioses are limited but growing (8,9).

The endosymbionts which colonize the cytoplasm of hindgut protists of *Reticulitermes* termites are from several bacterial taxa and vary between protist and termite species. These include *Endomicrobium* (10), and ‘*Candidatus* Ancillula’ (11). In addition, the nuclei of some hindgut protist species are colonized by *Verrucomicrobia* (12). Genome analysis of some of these endosymbionts, and some ectosymbionts, suggests that there is convergent evolution for these symbionts to support their unicellular host by synthesizing nutrients absent in the termites’ diet (11,13,14). Regarding *Endomicrobium*, previous studies have investigated both their population structure and transmission in *Trichonympha* spp. protists. Across various *Trichonympha* spp. these endosymbionts share rRNA gene phylogenies that are congruent with their host (15), are composed of a single phylotype in each host species, and are not shared across different *Trichonympha* spp. (16). These data suggest that these endosymbionts are vertically acquired.

The associations between termite protists and their symbiotic bacteria are only beginning to be understood. For example, in *Reticulitermes* spp. different protist species associate with *Treponema* from the same phylogenetic clusters, but the composition and fidelity of those associations are often not resolved beyond those broad phylogenetic groups. Furthermore, assessing the diversity of bacteria which associate with termite protists has been challenging since these protists are not yet cultivated, and are hard to isolate from their termite hosts. Previous studies have overcome these challenges by either using samples which consisted of pooled protist cells (3) or single-cell samples in which whole genome amplification (WGA) was performed (16). These methods have yielded novel information, but with recent advances in high-throughput amplicon sequencing and analysis we wondered if we could assay single-cells without the need to pool individuals or perform WGA.

Here, we used a method in which single protist cells, isolated from the hindgut of *R. flavipes*, served as a template for high-throughput amplicon sequencing of the hypervariable V4 region of bacterial symbiont 16S rRNA genes. We also co-amplified the 18S rRNA gene of some of the protist cells. Using these methods, the bacterial communities associated with single protist cells were investigated at single nucleotide resolution and high coverage using ASV methodologies. We selected six different protist species, four from the genus *Dinenympha* (Oxymonadida), *Pyrsonympha vertens* (Oxymonadida), and *Trichonympha agilis* (Parabasalia). We characterized the microbial communities associated with these protists which vary in their evolutionary relatedness and niches they occupy in the host. The ASV data also allowed us to investigate if bacterial communities associated with specific protist species varied across termite colonies (17).

We found that different protists species share some ectosymbiotic ASVs. This suggested that these particular symbionts may be cosmopolitan and perhaps horizontally acquired by their protist hosts. Horizontal transmission in these cases would result from the binding of ectosymbionts from the termite’s hindgut to new protist hosts. To test this hypothesis, we developed an *in vitro* fluorescence assay which allowed us to detect the horizontal acquisition of bacteria by protists. We show that the horizontal acquisition of these bacteria required active biological processes and the symbionts exhibited preferential spatial binding to their host cells in some cases.

## Materials and methods

### Termite collection, maintenance, and identification

*R. flavipes* termites were collected from three geographical sites using cardboard traps placed under logs for 2 to 4 weeks at the UConn Campus at Storrs, Connecticut (Geosite A/Colony A: Longitude −72.262216, Latitude 41.806543); in Granby, Connecticut (Geosite B/ Colony B: Longitude −72.789053, Latitude 41.999316) and in Mansfield Center, Connecticut (Geosite C/Colony C: Longitude −72.216138, Latitude 41.759949). Termites were removed from traps and maintained in plastic containers with moistened sterile sand and spruce wood. Species identity of the termites was verified to be *R. flavipes* by solider caste morphology (18), the presence of *Dinenympha gracilis* in the hindguts of worker termites (19,20), and sequencing of the cytochrome oxidase II gene (S3 Figure) with primers A-tLEU: 5’-ATGGCAGATTAGTGCAATGG-3’ (forward) and B-tLys: 5’-GTTTAAGAGACCAGTACTTG-3’ (reverse)(21). Only individuals of the worker caste were chosen for experiments described here.

### Amplification and sequencing of protist and bacterial SSU rRNA genes

Samples consisting of single protist cells were prepared from termites in an anaerobic chamber with atmospheric content of CO_2_ 5.5%, H_2_ 5.5%, and N_2_ 89%. Each hindgut was dissected and ruptured in 500 μl of Trager’s solution U (TU) (22). Hindgut contents were washed by centrifugation at 3,000 rpm for 90 seconds in an Eppendorf microcentrifuge and then resuspended in 500 μl of TU. This was done for a total of three washes. This washed cell suspension was then diluted 10-fold in TU. A 1 μl aliquot of the washed and diluted cell suspension was added to a 9 μl droplet on a glass slide treated with RNase AWAY® Reagent (Life Technologies) and UV light. Individual protist cells were isolated using a micromanipulator (Eppendorf CellTram® Vario) equipped with a hand-drawn glass capillary. Individual cells were washed three times in 10 μl droplets of TU via micromanipulation, transferring approximately 0.1 μl each time, and finally placed in 10μl molecular grade TE Buffer, and frozen at −20°C. If each gut sample contained 10^7^ bacteria, we estimate the final isolated protist cell should have less than one non-symbiotic bacterium from the original gut suspension. In total, we collected and analyzed 57 protist cells from 13 worker termites which were sampled from colonies at three geographical locations (Geosites A, B, and C).

Frozen protist cells served as templates for PCR reactions in which the 18S rRNA gene of the protist host as well as the V4 hypervariable region of the 16S rRNA gene of bacteria were co-amplified and sequenced. PCR reactions consisted of Phusion® High-fidelity polymerase (1 unit), HF buffer, dNTPs (200 μM), dimethyl sulfoxide (DMSO) (3%), 0.3 μM of each 18S primer (Euk19f, 5’-AYYTGGTTGATYCTGCCA-3’ and Euk1772r; 5’-CBGCAGGTTCACCTAC-3’) (23), 0.2 μM each of V4 16S primers (515f; 5’-GTGCCAGCMGCCGCGGTAA-3’ and 806r; 5’-GGACTACHVGGGTWTCTAAT-3’) (24), and a single protist cell in a final reaction volume of 50 μl. PCR conditions were as follows: Initial denaturation was at 94°C for 3 minutes followed by 35 cycles of 94°C for 45 seconds, 50°C for 60 seconds, 72°C for 2 minutes. Final extension was at 72°C for 10 minutes (17). For *P. vertens* and *D. gracilis* 18S primers 18SFU; 5’-ATGCTTGTCTCAAAGGRYTAAGCCATGC-3’ and 18SRU; 5’-CWGGTTCACCWACGGAAACCTTGTTACG-3’ were used (25).

PCR products were separated and visualized on a 1% agarose gel, gel-purified and quantified using Qubit™ Flourometric quantitation (ThermoFisher Scientific). Barcoded V4 16S rRNA gene amplicons were pooled at 4nM equimolar concentrations and sequenced on an Illumina Miseq (17). Amplicon libraries were generated using both single (24 samples) and dual (58 samples) indexed primer sets. Negative controls consisting of TU, TE, and protist-free technical controls were amplified and sequenced in order to monitor contamination from slides, buffers, kits and other reagents. 18S rRNA gene amplicons were cloned using the pGEM®-T Easy Vector System (Promega) and sequenced by Sanger sequencing. If needed, additional isolated protist cells were used in 18S rRNA gene-only PCR reactions and the amplicons were cloned and sequenced as described above.

### V4 16S amplicon analysis using ASV inference

V4 16S gene reads were analyzed using a single workflow in R. Raw V4 amplicon reads were initially processed using the R package DADA2 (26). Reads were quality trimmed, filtered, merged, and exact biological sequences (ASVs) were determined using the Divisive Amplicon Denoising Algorithm (DADA) (26). This allows for determining the true biological sequence of V4 amplicons by applying the error rates observed within samples (26, 27). Taxonomy was then assigned to ASVs in DADA2 using the SILVA rRNA gene database Version 132 (28). Reagent and other contaminants were then identified and removed using the R package decontam using the prevalence method with the threshold set to 0.35 (29). This allowed identification of contaminating ASVs by assessing the prevalence of ASVs in negative control samples compared to protist samples (29). Using taxonomic assignments, ASVs were then filtered based on their ecologically plausibility such that only previously reported protist-associated bacterial taxa remained using the R package Phyloseq (30). We then removed ASVs that were not found in at least three of the protist samples and at least 1 x 10^-5^ in their relative abundance. These filtering parameters are consistent with recent studies that identified contaminants in low-biomass samples (31). Exact filtering commands are provided within the amplicon workflow in S2 File. Ordination analysis, α-diversity (Observed, Chao1, and Shannon), and β-diversity analysis (Bray-Curtis metric) were done using Phyloseq and Vegan (32) and data were plotted using the R package ggplot2 (33). For ordination analysis, a NMDS plot was generated using the Bray-Curtis metric and ellipses were generated for each protist genus per colony, representing the 95% confidence intervals. Statistical significance between α-diversity measurements was assessed using the Mann-Whitney test in GraphPad Prism (version 8). Adonis which is part of the Vegan R package was used to perform PERMANOVA analyses. Rarefaction analysis was performed using the “ggrare” function of ggplot2.

### Phylogenetic analysis of SSU rRNA genes

ASVs corresponding to *Endomicrobium* were aligned to V4 16S rRNA gene reference sequences using MUSCLE (34). Phylogenetic trees were generated using a Maximum likelihood (ML) method with the program IQ-TREE with model testing (35). The 18S rRNA genes obtained by this study were also aligned to reference sequences using MUSCLE and a phylogenetic tree was made using IQ-TREE with model testing.

### Scanning electron microscopy

Protist cells were collected by low spin centrifugation as described above and fixed in 2% glutaraldehyde in TU (pH 7) for 1 hour at RT in an anaerobic chamber. The samples were deposited onto poly-L-lysine coated silicon wafer chips (Prod No. 16008, Ted Pella Inc.), washed with 80 mM Na cacodylate buffer (pH 7), and post-fixed in 2% osmium tetroxide at RT for 1 hour. The cells were rinsed twice for 5 minutes in distilled water then dehydrated in serial concentrations of ethanol (30%, 50%, 70%, 95%, 100%, 5 min each) and critical point dried (931GL, Tousimis). Samples were then mounted on SEM stubs using silver paint, sputter coated with palladium (E5100, Polaron) and examined using a scanning electron microscope (Nova NanoSEM 450, FEI).

### Live fluorescent symbiont transmission assays

For all live-transmission assays, experiments were carried out in an anaerobic chamber with gas composition as described above. Hindguts were dissected from termites, ruptured with sterile forceps, and their contents were collected in anaerobic TU buffer containing resazurin (1μg/ml), sodium thioglycolate (0.5g/L), and sodium bicarbonate (0.1M) pH 8.0 (Pedro et al., 2004). Samples were then fractionated by low spin centrifugation (3,000 rpm for 90 seconds) to separate protists and their symbionts and from bacteria which were unattached to protists. Bacteria and protists were treated separately in order to minimize damage to protists by the high centrifugation forces needed to pellet bacteria. Each fraction was then washed three times in buffer by centrifugation at either 3,000 rpm (for protist fraction) or 13,000 rpm (for bacterial fraction) for 90 seconds. The washed fractions were then split into two equal volumes and stained with either Texas Red®-X succinimidyl ester (TRSE, Molecular Probes™) or AlexaFlour Green 488 succinimidyl ester (SE488, Molecular Probes™) at 10μg/ml for 1 hour at room temperature (RT) in the dark in the anaerobic chamber. Covalent dye conjugation was done per manufacturer’s instructions. Following dye conjugation, cells were washed three times in TU with reduced glutathione serving as a stop reagent for the amine reactive dyes. Protist and bacterial fractions were combined to produce two samples: red protists plus red free-living bacteria and green protists plus green free-living bacteria.

To assay for symbiont acquisition by protists, half of each sample, red and green, were combined and monitored for the horizontal acquisition of new bacteria which was evident by heterogeneity in fluorescent signals of bacteria on individual protists. Subsamples were taken at 0, 3, 15, and 20 hours, fixed with 2% formaldehyde, and viewed using a Nikon TE300 Eclipse microscope. Alternatively, fixed samples were mounted in ProLong™ Diamond Antifade Mount (ThermoFisher) and imaged using a Nikon A1R Spectral Confocal microscope. To determine specificity of binding locations on protist hosts the positions of newly acquired ectosymbionts were counted along the length of two *Dinenympha* species (*D.* species II and *D. fimbriata*) and correlations in binding towards the anterior or posterior cell poles were tested using the Pearson R correlation test in GraphPad Prism (version 8). To test if symbiont acquisition required biologically active processes, this assay was repeated with the addition of either tetracycline 10μg/ml or cycloheximide 10μg/ml to each sample 1 hour prior to the start of the assay and compared to a no-treatment control. In addition, anaerobic symbionts were killed by exposure to atmospheric oxygen, labeled with propidium iodide (PI), and mixed with live cells to assay for the binding of dead bacteria to live protist hosts.

The fluorescent assay was then used to investigate whether ectosymbionts could come from the free-living (unattached) pool of bacteria. Hindgut contents were fractionated into a free-living bacterial fraction and a protist-cell fraction as described above and each was stained red with TRSE. These were added separately to a green SE-488 stained protist-cell fraction and incubated in an anaerobic chamber as described above. Samples were taken at 15 hours post the start of the assay, fixed, and viewed as described above.

## Results

### Termite mitochondrial cytochrome oxidase II gene sequencing and phylogenetic analysis

The mitochondrial cytochrome oxidase II gene was sequenced from worker termites from all three colonies that were used in this study. Phylogenetic analysis indicates that these sequences grouped with publicly available references of *R. flavipes* from other studies (S3 Figure).

### Morphological and phylogenetic diversity of hindgut protists

The protists used in this study were investigated using both light microscopy, SEM and 18S rDNA sequencing. These data indicated that these protists belonged to six different species: *T. agilis*, *P. vertens*, *D. gracilis*, *D. fimbriata*, and two uncharacterized *Dinenympha* species (I & II). We obtained near-full length or partial (>1 kb) 18S rRNA genes sequences from individual protist cells, aligned them to references sequences, and reconstructed their phylogeny using IQ-TREE. The previously undescribed organisms referred to here as “*Dinenympha* species I & II” clustered with other *Dinenympha.* Based on their differing 18S genes and their differing morphologies, we refer to *D. gracilis*, *D. fimbriata*, *D.* species I and *D.* species II as separate species in this paper (Figure 1). In depth studies of the gene evolution will be required to determine the exact relationships of these organisms to each other. DIC micrographs of representative morphotypes of each protist species used in this study are provided in Figure 1.

**Figure 1.**
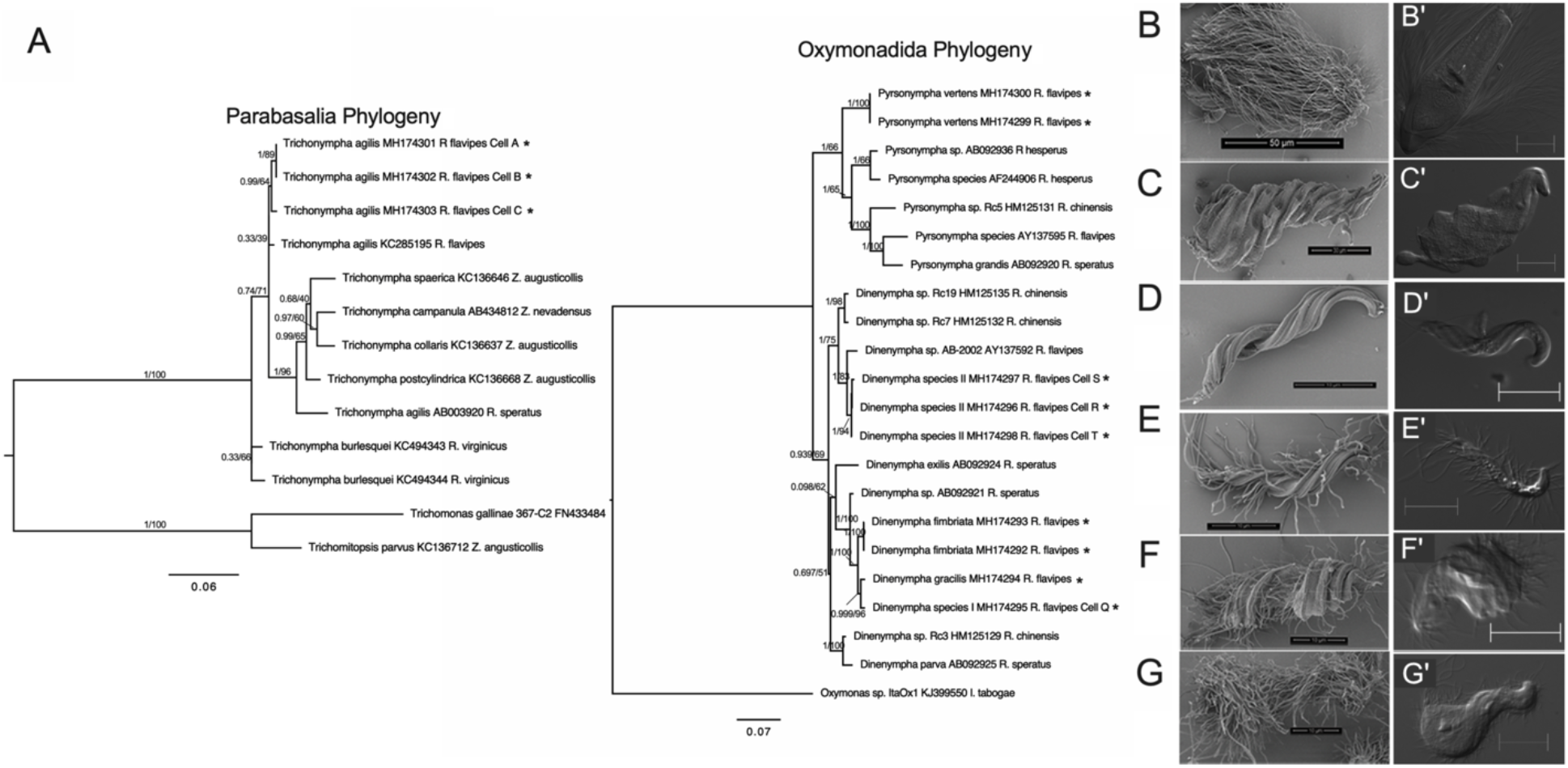
Phylogenetic and morphological diversity of hindgut protist species from *R. flavipes*. (A) Maximum likelihood (ML) phylogenetic trees of 18S rRNA genes from single protists cells. Parabasalia ML tree (left) was generated using the program IQ-Tree with substitution model GTR+I+G4 while the Oxymonadida ML tree (right) was generated using the substitution model TN+I+G4. Four 18S rRNA genes from single protist clustered to known references sequences (*D. fimbriata*, *D. gracilis*, *P. vertens*, and *T. agilis*). Other previously undescribed protists, *D*. species I and *D*. species II, clustered within the genus *Dinenympha*. Taxa marked by (*) represents sequences obtained by this study. Sequences from cells which are designated by a letter represent individuals in which the 18S rRNA gene was co-amplified with the bacterial V4 16S rRNA gene. Tip labels include the protist name, accession number, and host termite species. Support values represent the Bayesian posterior probability and Bootstrap support values respectively. Scanning electron micrographs, and DIC images of representative individuals of each protist species ((B, B’) *T. agilis*, (C, C’) *P. vertens*, (D, D’) *D. gracilis*, (E, E’) *D. fimbriata*, (F, F’) *D*. species I and (G, G’) *D*. species II. Electron micrograph scale bars represent 50μm (B), 30μm (C), and 10μm (D – G), all DIC scale bars are 20μm.

### ASV composition of protist-associated bacteria

High-throughput amplicon analysis of the V4 region of the bacterial 16S rRNA gene recovered 102 bacterial ASVs from single protist cells spanning six different protist species. The most diverse bacterial taxon was the genus *Treponema* which accounted for 66 of the 102 ASVs (Figure 2A). Other taxa included the genera *Endomicrobium* (7 ASVs) and the phyla Verrucomicrobia (3 ASVs), Bacteroidetes (17 ASVs), Proteobacteria (7 ASVs) Margulisbacteria (1 ASV) and Synergistetes (1 ASV) (Figure 2A).

**Figure 2.**
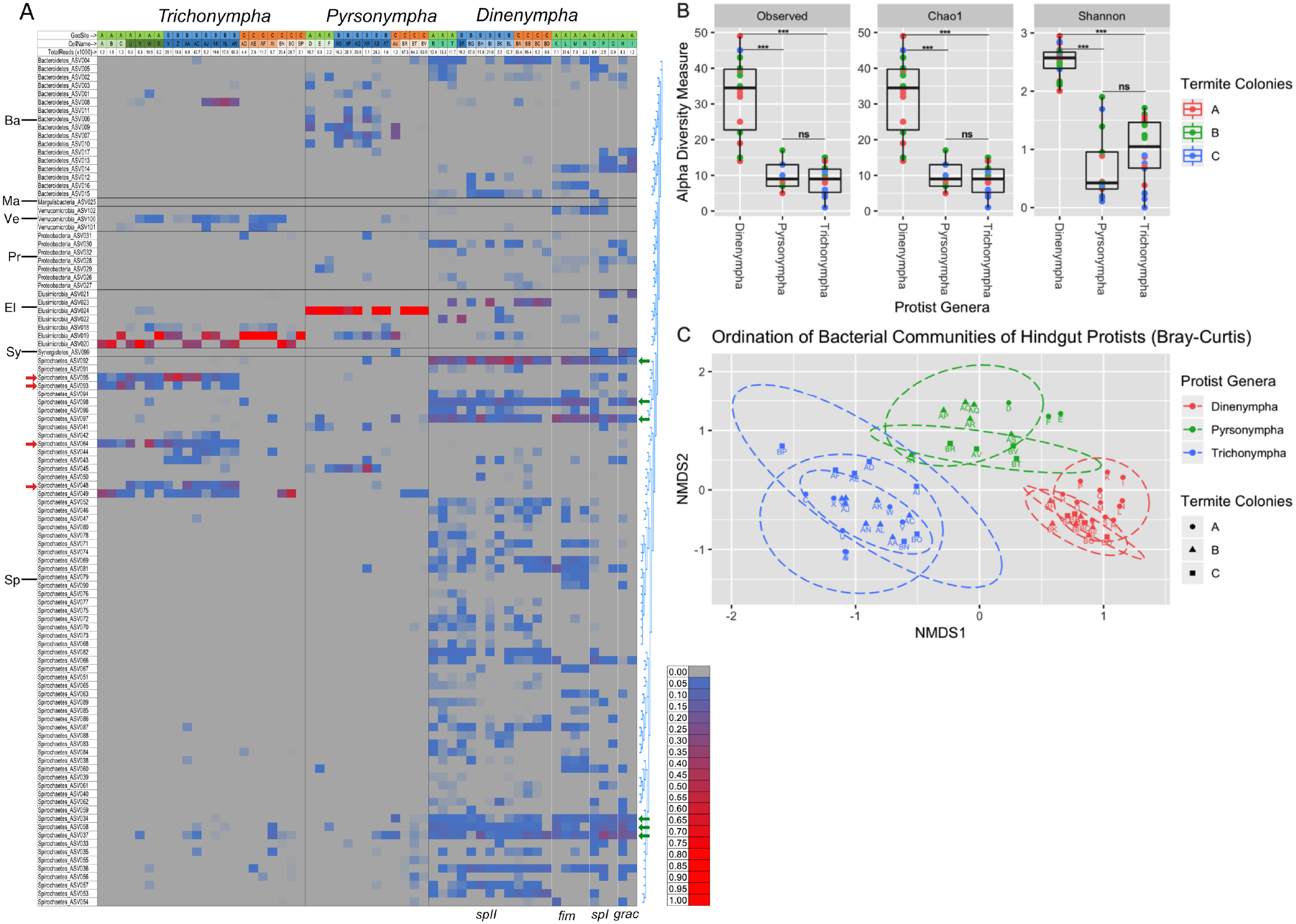
Diversity and distribution of protist-associated bacterial ASVs. (A) Heat map showing the distribution and relative abundance of bacterial ASVs (rows) which are grouped by similarity in the tree at right (generated by Muscle alignment and Jukes-Cantor Neighbor Joining as implemented in Geneious v9) and at the Phylum level on the left: Ba=Bacteroidetes, Ma=Margulisbacteria, Ve=Verrucomicrobia, Pr= Proteobacteria, El=Elusimicrobia, Sy=Synergistetes, Sp=Spirochaetes. Individual protist cells are in the grey columns, geosites, cell_ID and total reads per sample are indicated at the top. Red arrows indicate Spirochaetes ASVs which were present on *Trichonympha* hosts from colonies A and B but absent on those from colony C. Green arrows indicate Spirochaetes ASVs which were present on all four species of *Dinenympha* hosts. (B) α-diversity measurements (Observed, Choa1, and Shannon) of each of the three protist genera (*Dinenympha*, *Pyrsonympha*, and *Trichonympha*). Box plots represent the 25th and 75th percentiles with the line representing the median and whiskers representing the largest and smallest values, respectively. Statistically significance was calculated using Mann-Whitney test (***, p< 0.0001, ns= no significant difference). (C) Bray-Curtis β-diversity ordination plot of the bacterial ASVs of protist cells. Each point represents the bacterial community of a single protist cell where the color represents the protist genus, shape indicates the termite colony collection site, and the dashed ellipses represents the 95% confidence intervals for each protist genus per termite colony collection site. Points are labeled by a letter which represents protist cell’s identification. Green and red arrows are explained in the Results section.

To determine if there were any differences between the communities of bacterial symbionts across different protist genera, α-diversity and β-diversity analyses were conducted using the Phyloseq and Vegan R packages. Despite being smaller in size, *Dinenympha* species hosts were associated to a more diverse bacterial community compared to both *Trichonympha* (Mann-Whitney test Observed, Chao1, and Shannon α-diversity p < 0.0001) and *Pyrsonympha* (Mann-Whitney test Observed, Chao1, and Shannon β-diversity p < 0.0001) protists (Figure 2B). Rarefaction analysis indicated that each protist sample was sequenced at sufficient coverage to determine their symbiont’s taxonomic diversity (S4 Figure). This was performed with and without sequences corresponding to *Endomicrobium* (endosymbionts) to ensure that less abundant taxa were also sequenced at sufficient coverage. Thus, it is not likely that differences in community diversity between protist species were artifacts of inadequate sequence depth coverage.

The protist-associated communities grouped together according to their protist hosts using the Bray-Curtis metric to perform β-diversity ordination (Figure 2C). The communities were significantly different from one another and were influenced by the genus of their host protist (PERMANOVA Bray-Curtis *f*=24.188, *R^2^*=0.47253, *p*=0.001) and to lesser extent, the collection site from which they were originally isolated (PERMANOVA Bray-Curtis *f*=1.8819, *R^2^*=0.06516, *p*=0.041). The effect of termite geographical collection site was further investigated within each group of protist hosts. For *Trichonympha* hosts, the bacterial communities were heterogeneous with respect to termite collection site (PERMANOVA Bray-Curtis *f*=3.3298, *R^2^*=0.25954, *p*=0.019). This heterogeneity can be most easily seen for Spirochaetes ASVs 95, 93, 64 and 48 (Fig 2A, red arrows), which were all absent in *Trichonympha* cells from termites collected in geosite C, but were present in *Trichonympha* from geosites A and B. Bacterial communities associated with *Dinenympha* were also heterogeneous with respect to termite collection site (PERMANOVA Bray-Curtis *f*=3.1712, *R^2^*=0.25027, *p*=0.001) however, the bacterial communities of *Pyrsonympha* hosts were homogenous across different termite colonies (PERMANOVA Bray-Curtis *f*=0.633226, *R^2^*=0.11226, *p*=0.677). Collectively, these data suggest that the bacterial communities of each protist genus are distinct from one another and can show some collection site differences for some bacteria that associate with some protist types.

Several ASVs, likely corresponding to ectosymbionts (Spirochaetes), were observed to be present across almost all of the *Dinenympha* species including *D. gracilis* which has very few visible ectosymbionts when observed by microscopy. (Figure 1 and Figure 2A). ASVs 92, 97, 98, 34, 58 and 37 (Fig 2A, green arrows) appear to be associated with all four species at relatively high levels: 5% - 10% in *Dinenympha* species II and *D. fimbriata*, 5% - 30% in *Dinenympha* species I and 3% - 20% in *D. gracilis*. ASV36 was found to be associated with all four *Dinenympha* species though at lower levels: 0.5% - 5%. In addition, these ASVs were also associated with some of the *Trichonympha* and *Pyrsonympha* protist cells, but only sporadically.

### Phylogenetic diversity of protist-associated *Endomicrobium*

Previous studies concluded that the *Endomicrobium* endosymbionts demonstrate co-speciation with their protist host as inferred by congruent phylogenies (37). We investigated which of the *Endomicrobium* (Elusimicrobia) ASVs demonstrated that same congruence. This was possible since we were able to co-amplify and sequence some of the 18S rRNA genes of protist cells used in our high-throughput amplicon study. The results from these phylogenetic analyses suggested that five out of the seven *Endomicrobium* ASVs shared congruent evolutionary histories with their protist hosts while ASV18 and ASV21 did not (S5 Figure). ASV21 was associated at low abundance with only three *Dinenympha* cells and may be a contaminant. ASV18, however was found associated, at low abundance (generally <1%) with 11 *Trichonympha* cells (50%), and may be a contaminant, or a low-abundance symbiont of these protists. The remaining five *Endomicrobium* ASVs clustered into three distinct groups, each associating with either *Trichonympha* (ASV19 and ASV20), *Pyrsonympha* (ASV24), or *Dinenympha* (ASV22 and ASV23). The two *Endomicrobium* ASVs that were associated with *Trichonympha,* hosts differed from one another by one base pair and did not appear to co-colonize individual *Trichonympha* cells. This trend was also true for the two *Endomicrobium* ASVs which associated with *Dinenympha* species II (Figure 2A). We did detect some instances in which some of the *Endomicrobium* sequences from one host were present in other protist samples. For example, one of the *Endomicrobium* ASVs from *Trichonympha* hosts (ASV19) was found in several *Pyrsonympha* cells. This ASV was present at high relative levels in those *Pyrsonympha* cells that did not have high total read counts and may represent contamination from lysed *Trichonympha* cells (Figure 2A).

### Horizontal acquisition of labeled bacteria

The ASV analysis, outlined above, suggested that some ectosymbiont types (Spirochaetes) are found associated with multiple protist species. This could occur through the horizontal transmission of ectosymbionts. An *in vitro* fluorescence-based assay was developed to test this possibility. Protists and bacteria from the hindgut of *R. flavipes* were covalently stained with either TRSE (red fluorescence) or SE488 (green fluorescence), mixed together, and the acquisition of new ectosymbionts was assayed over time. Since protists began the experiment with ectosymbiotic communities that were homogeneous in their fluorescent label, newly acquired bacteria could be detected if individual protists acquired bacteria of a different color over time. Acquisition observed in this assay should represent only half of the total transfer events since we could not distinguish newly acquired bacteria that were the same color as the majority of the cells on the host. Over time, we observed that protist hosts including *T. agilis, D. fimbriata* and *D.* species II acquired new bacteria (Figure 3) that appeared to be attached to host’s plasma membrane (Figures 3E, 3J, and 3O), not merely entangled in flagella or entangled with other bacterial cells. Such events were never observed in non-mixed control samples indicating that what appeared to be acquired bacteria were not bacteria with unusual autofluorescence properties (S6 Figure). We used our fluorescence assay to determine if newly attached bacteria could transfer to protists from just the pool of free-living (unattached) bacteria, or just from other protist cells. (Figure 3P). In these experiments there was no significant difference between the percentage of protist cells that acquired new ectosymbionts from the free-living or protist-associated fractions (Two-tailed T-test, t = 1.054, df = 2, p = 0.402).

**Figure 3.**
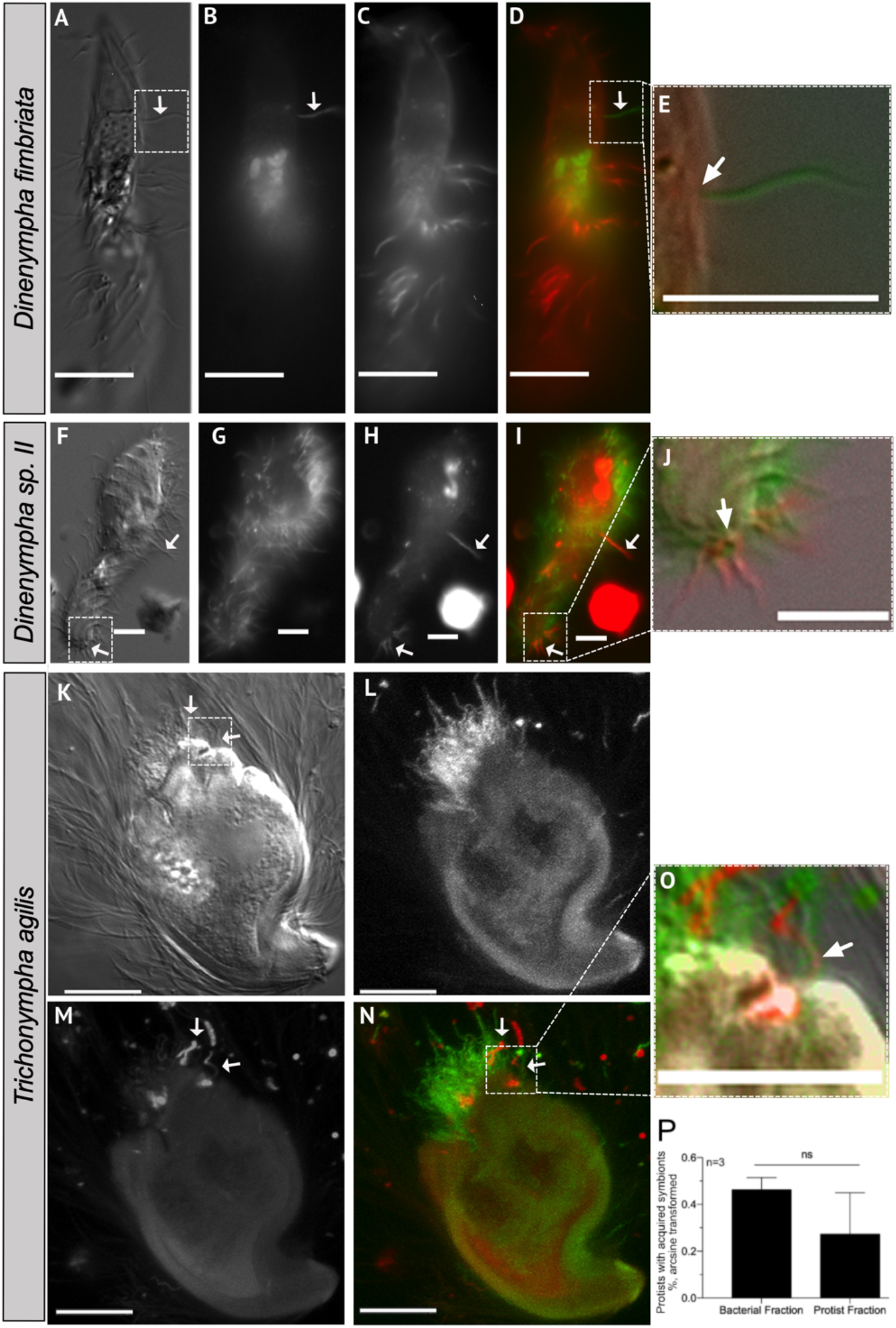
Horizontal acquisition of ectosymbionts by different protist species. DIC and fluorescence micrographs of hindgut protist and their ectosymbionts stained with either TRSE (shown red) or SE-488 (shown green) at Time=12 hours of fluorescent acquisition assay. Overtime several different protist species including (A-E) *D. fimbriata*, (F - J) *D*. species II, and (K – O) *T. agilis* acquired new ectosymbionts. Micrographs are arranged from left to right as DIC, SE488, TRSE, and merged (SE488 and TRSE) for each protist. Fluorescence micrographs L – M are maximum intensity Z-projections. Micrographs E, J, and O represent areas which are zoomed in to demonstrate that the ectosymbionts are visibly attached to the host cell’s plasma membrane. Arrows point to horizontally acquired ectosymbionts and scale bars represent 10μm. (P) The average percent (arcsine transformed) of protists that acquired new bacteria from either the protist or bacterial cell fractions of termite hindguts at Time=15 hours of the assays. Error bars represent the standard error of the mean and n= 3 independent experiments.

In well characterized symbioses in which symbionts are horizontally transmitted, several active biological processes are involved. These include changes in the gene expression of the symbiont so that it can properly recognize and occupy its niche on, or in, its host (38,39). To determine if bacteria acquisition by hindgut protists required active processes, we tested whether inhibiting protein synthesis affected the acquisition assay. The assay was done with the addition of either tetracycline or cycloheximide and compared to a no-treatment control. Tetracycline, which inhibits prokaryotic translation, was chosen because of previous reports that termite-associated Spirochaetes and Bacteriodetes are sensitive to that antibiotic (40). Cycloheximide has been used to target protein synthesis across different protist taxa (41,42). Over time, samples which were treated with tetracycline had significantly fewer protists that acquired new ectosymbionts compared to the no-treatment control (Two-tailed T-test, Time = 20 hours, t = 5.278, df = 3, p = 0.0133) (Figure 4A). Other control experiments showed that bacteria were not killed by tetracycline over the 20 course of the experiment (not shown) These data indicated that inhibiting protein synthesis in the bacteria may have affected their ability to be horizontally acquired by their protist hosts. However, we cannot rule out that tetracycline may have affected host cells and lowered transmission rates. Cycloheximide treated samples were not significantly different from the no-treatment control. However, it may still be the case that protein synthesis by the protist host is required for acquisition if the rate of protein turnover is slow and our 20-hour assay was not long enough time to detect an effect, or if these protists are not sensitive to cycloheximide.

**Figure 4.**
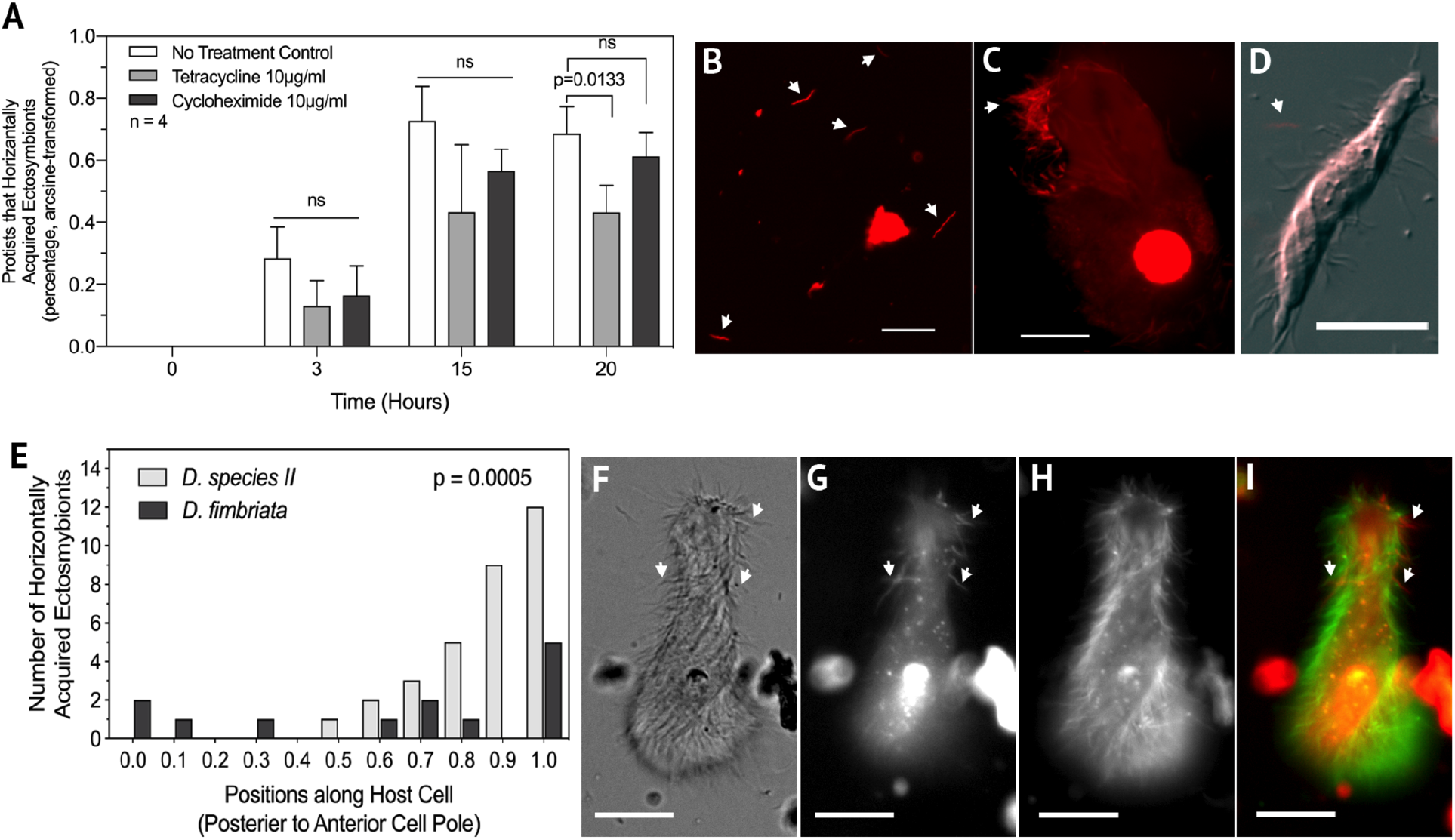
Horizontal transmission of ectosymbionts involves active processes and is non-random. (A) The addition of tetracycline significantly lowered the percentage of protists that acquired at least one new ectosymbiont (Two-tailed T-test, Time = 20 hours, t = 5.278, df = 3, p = 0.0133, n = 4 independent experiments, and FDR Statistics available in S1 File) while the addition of cycloheximide had no significant effect. (B - D) Micrographs of propidium iodide stained cells. Exposing hindgut contents to O2 killed hindgut bacteria (B and C) which did not bind to live protist cells (D) (arrows point to O2 killed bacteria). (E) Significantly more ectosymbionts (Pearson’s R, p=0.0005) bound towards the anterior cell pole compared to the posterior cell pole on *D*. species II however, this binding characteristic was not seen in other *Dinenympha* species. (F - I) Fluorescence and DIC micrograph of *D*. species II stained with amine reactive dyes (G TRSE, H SE488), showing increased binding of new ectosymbionts (arrows) toward the anterior cell pole. Scale bars represent 10μm.

In addition, protists and bacteria were exposed to atmospheric oxygen for several hours. We confirmed that oxygen killed both ectosymbiont and free-living bacteria by labeling with propidium iodide (PI) which labels cells which have died (43) (Figure 4B and 4C). These PI-labeled cells were then added to live samples to assay for the binding of dead bacteria to live protist hosts. In these experiments, we did not observe the binding of dead ectosymbionts to live protist cells (n = 4 independent experiments) (Figure 4D). Thus, acquisition of bacteria in this assay required live partners and active translation.

We noticed that most of the newly acquired bacteria appeared to bind to the anterior end of *Dinenympha* species II. To determine if this was true, or if binding was random, newly attached bacteria were counted along the length of this protist species. The resulting data indicated that newly acquired bacteria bound more frequently towards the anterior cell pole of *D*. species II (Pearson’s R p=0.0005) (Figure 4E-4I) than the posterior cell pole. This increase in frequency at one cell pole compared to the other was not observed in other *Dinenympha* species (Figure 4E). This suggested that in *Dinenympha* species II there was a preferred region for the addition of newly acquired bacteria.

## Discussion

In this study, we found that the associations between *R. flavipes* hindgut protists and their symbiotic bacteria exhibited specificity in host range and community structures. We also found that *Dinenympha* species, as a group, associated with a more diverse community of bacteria compared to the larger protist species *P. vertens* and *T. agilis* (Figure 2B). Since most of the ASVs associated with these *Dinenympha* were *Treponema* (Figure 2A), which are considered to be mainly ectosymbionts, these data suggested that it was the ectosymbiont communities of these *Dinenympha* species that were responsible for the increased diversity. These ectosymbiont communities included ASVs that were shared among some *Dinenympha* species which led to the hypothesis that some ectosymbionts may be horizontally acquired or transferred.

In general, the bacterial communities of the protist species studied were distinct from one another (Figure 2C) as most bacterial ASVs were exclusive to certain protist hosts or classes of protist hosts (Figure 2A). However, the individual associations between symbionts and protist species exhibited varying levels of fidelity. For example, even though many *Treponema* ASVs were shared between *Dinenympha* species, only a few were found to occasionally be associated with *Trichonympha* or *Pyrsonympha* (Figure 2A). The converse was also true: *Treponema* ASVs associated with *Trichonympha* or *Pyrsonympha* were only occasionally associated with *Dinenympha* cells. This suggested that there may be mechanisms that result in, or ensure, specificity between bacterial symbionts and a range of related protist hosts. The *Treponema* diversity associated with *Dinenympha* hosts may be due to common features such as attachment factors which are shared among protist hosts who are closely related and co-inhabit the same termite hindgut.

These protist-bacteria associations were relatively conserved across different termite colonies (Geosites), however some of the variation within these groups of protists was due to collection site effects (Figure 2C). This results of colony-level variation of protist-bacterial associations complements previous reports of variation in the microbial communities of termite guts (17,44). This repeated observation across different studies highlights the importance of using colony replication in termite microbiome research.

The ASV analysis provided a high-resolution characterization of the community structures of the endosymbionts of hindgut protists. For example, individual *T. agilis* cells differed with respect to which *Endomicrobium* ASV they were associated. Two *T. agilis* cells (cells A and B) which shared the same *Endomicrobium* ASV were more similar to one another in their 18S rRNA gene sequence than they were to a third cell (cell C, Figure 1A) which associated with a different *Endomicrobium* ASV. Even though these two *Endomicrobium* ASVs differed by only one base pair in the V4 region genome assembly and analysis shows that they are distinct genomovars or strains (unpublished data). Previous studies have demonstrated the possibility that what was thought to be a single species of *T. agilis* in *R. flavipes* is likely more than one species (45). Our data, showing two ASVs with related, but distinct genomes, that colonize *Trichonympha* cells with different 18s rRNA gene sequences support that possibility.

The ectosymbiont communities of hindgut protists in *R. flavipes* may not always be static as demonstrated by the fact that we observed that protists could acquire new bacteria on their surfaces over time. This horizontal acquisition of bacteria was likely to be an active process because transfer was lowered in the presence of tetracycline (Figure 4A). This less effective transfer may have been caused by inhibition of bacterial translation of proteins directly needed for binding, or due to indirect effects. For example, tetracycline treatment may have affected motility, chemotaxis, or protist functions as was shown in an earlier study (40). The inability to cultivate these members currently limits their experimental tractability and at this time we are unable to determine the exact mechanisms of how tetracycline lowers symbiont transmission. Oxygen-killed hindgut bacteria were never observed to attach to live protist hosts indicating that the horizontal acquisition seen in our experiments is likely active and not due to non-specific binding or entanglement. The observation that ectosymbionts bound preferentially to the anterior cell pole of *D*. species II suggested that there was spatial specificity to the process (Figure 4E-4I). This spatial specificity was not observed with other *Dinenympha* species. The cause of this spatial preference is not known, but may be the result of new cell membrane, or binding structures, being added to the host at the anterior pole. Because ectosymbionts are found at other locations along the cell, ectosymbionts may be transported to these sites following binding at the anterior pole. Spatial specificity of ectosymbiotic *Treponema* on the surface of termite protists has been observed in a previous study using phylotype-specific fluorescent in situ hybridization (FISH) probes (3).

Protists from *R. flavipes* cannot yet be cultured, and this resulted in some limitations in the ectosymbiont acquisition assays. After 20 hours, most protist cells have died and lysed during the *in vitro* experiments. This limited the time over which the assay was conducted. Because of this, we could not determine if ectosymbionts could also be vertically transmitted during protist cell division. We have not witnessed actively dividing hindgut protists, but there is no evidence to suggest that they would have to shed their ectosymbionts prior to, or during, cell division. Thus, most surface-bound bacteria are likely partitioned between daughter cells during division.

Horizontal transmission of gut microbiota has occurred between different termite species and is thought to be a driving force in shaping their microbial communities (46). Here we extend the possibility of horizontal transmission occurring at the level of protist-bacterial symbioses within the guts of termite hosts. *R. flavipes* is a foraging wood-feeding termite which may cause them to have a greater influx of transient members and as a result have a greater bacterial species turnover compared to non-foraging (wood-dwelling) species (44). However, the termites used in this study were maintained under laboratory conditions on initially sterile sand and fed initially sterile wood which may reduce the introduction of transient members and select for a more stable gut community (44).

The results outlined above suggest that the acquisition of new bacteria by protists that we observed required live bacteria and was not due to non-specific interactions such as entanglement in protist flagella. However, we do not currently have direct evidence that the acquired bacteria are any of ectosymbionts characterized by our V4 analysis. They may be true symbionts, or they may be only transiently bound: staying for longer than our 20 hour assay, but perhaps not establishing a permanent association with a protist host. Using FISH with ASV-specific probes would help to resolve which bacteria were horizontally acquired. However, because as many as 30 or more *Treponema* ASVs associated with a single protist cell (as is the case with some *D*. species II cells), designing and testing that many FISH probes, and tying FISH results to a single, newly acquired bacteria would be impractical. Another approach that may work, but that would be technically challenging, would be to isolate single, transferred, bacteria from the surface of protists and characterize them by V4 analysis or single-cell sequencing.

Both transient and permanent binding to the surfaces of protists may aid in a bacterium’s ability to persist in the gut of a termite host by preventing them from being voided in termite’s fecal material by peristalsis. It may also give some members greater access to nutrients by allowing them to transverse the gut environment on a much larger and motile protist host. It has been proposed that at least some of the ectosymbiotic *Treponema* may be acetogens (4) such as the cultivated *Treponema primitia* strains (ZAS-1 and ZAS-2) which were isolated from *Zootermopsis angusticollis* and shown to carry out reductive acetogenesis (48). Associating with acetogens may be beneficial for protists which produce CO2 and H2 as end products from fermenting the polysaccharides found in wood since those products would be consumed during reductive acetogenesis (4).

Currently, the metabolic potential of many of these protist species and their symbiotic communities of bacteria remain unknown. It is generally accepted that at least some of the protist species are cellulolytic, which stems from the observations that some protists commonly contain wood particles in their cytoplasm (1) and some members, which were previously cultivated from another termite species, were shown to hydrolyze cellulose *in vitro* (49,50). There are recent reports that some ectosymbionts of Oxymonadida protists have the capacity to hydrolyze some of the polysaccharides found in wood (9,47). These include ‘*Ca*. Symbiothrix’ which are associated with *Dinenympha* species (5,9) as well as, several ectosymbionts of *S. strix* such as members of the Bacteroidetes and ‘*Ca*. Ordinivivax streblomastigis’ (47). It is unclear if the function of polysaccharide hydrolysis is limited to these symbionts or if it’s a general feature of the ectosymbionts of termite gut protists. Methods which use single protist cells, such as the amplicon analysis performed in this study, and single cell metagenomics (47,51) will help resolve the community structures, variation, and the potential functions of these protist-bacterial symbioses in termites.

The use of single protist cells as templates for high-throughput amplicon sequencing of the V4 region of their bacterial symbiont’s 16S rRNA gene combined with the high-resolution of the DADA2 analysis, provided a detailed survey of these bacterial communities. It is important to note that although we carefully isolated these single cells and washed them by micromanipulation, it is still plausible that either bacteria or DNA were carried over during isolation, and that some of the associations depicted in Figure 2A are artifacts of this kind. For example, we did detect some cases where *Endomicrobium* ASVs were found associated with the “wrong” protist types at a low prevalence and low abundance. Increasing the number of washing steps or more conservative filtering criteria may reduce or eliminate these issues in future studies. Our data supports the hypothesis that bacterial communities associated with different protist classes are distinct from one another, and that contamination and carry over was not great enough to obscure statistically significant differences in community structure (Figure 2).

Future studies could investigate the use of different combinations of universal primers sets to increase the success of co-amplifying the 18S rRNA genes from protist hosts along with the V4 (or other regions) of the bacterial 16S rRNA genes. Another way in which these single cell assays could be explored, or improved upon, would be to investigate the use of an 18S rRNA primer set which could amplify a variable region that would allow for high-throughput amplicon sequencing. This would remove the need to clone and sequence 18S amplicons by Sanger sequencing and likely yield higher coverage at a reduced cost. High-throughput amplicon sequencing to investigate protist diversity is already in use for termite protists, and other protist as well. (52).

## Supporting information

R script to repeat data analyses

## Author Contributions

MES and DJG designed experiments. MES performed experiments and data analysis. MES and DJG wrote the manuscript.

## Acknowledgments

We would like to thank Dr. Matthew Fullmer for discussions on phylogenic tree building. We would also like to thank Charles Bridges and Jaimie Micciulla for aiding in the collection and maintenance of termite colonies. SEM work was performed in part at the Bioscience Electron Microscopy Facility of the University of Connecticut. This research was funded by the National Science Foundation (NSF) division of Emerging Frontiers in Research and Innovation in Multicellular and Inter-kingdom Signaling. Award number 1137249 (DJG).

## Data Availability Statement

All 18S rRNA gene sequences derived from protists have been submitted to the National Center for Biotechnology Information (NCBI) Genbank under accession numbers MH174292 – MH174303 as well as the termite mitochondrial cytochrome oxidase II gene (accession numbers MH171305, MK950845, and MK950846). Sequences of the bacterial V4 16S rRNA gene amplicons have been submitted to the NCBI Sequence Read Archive (SRA) under BioProject PRJNA544517.

---

**S3 Figure.**
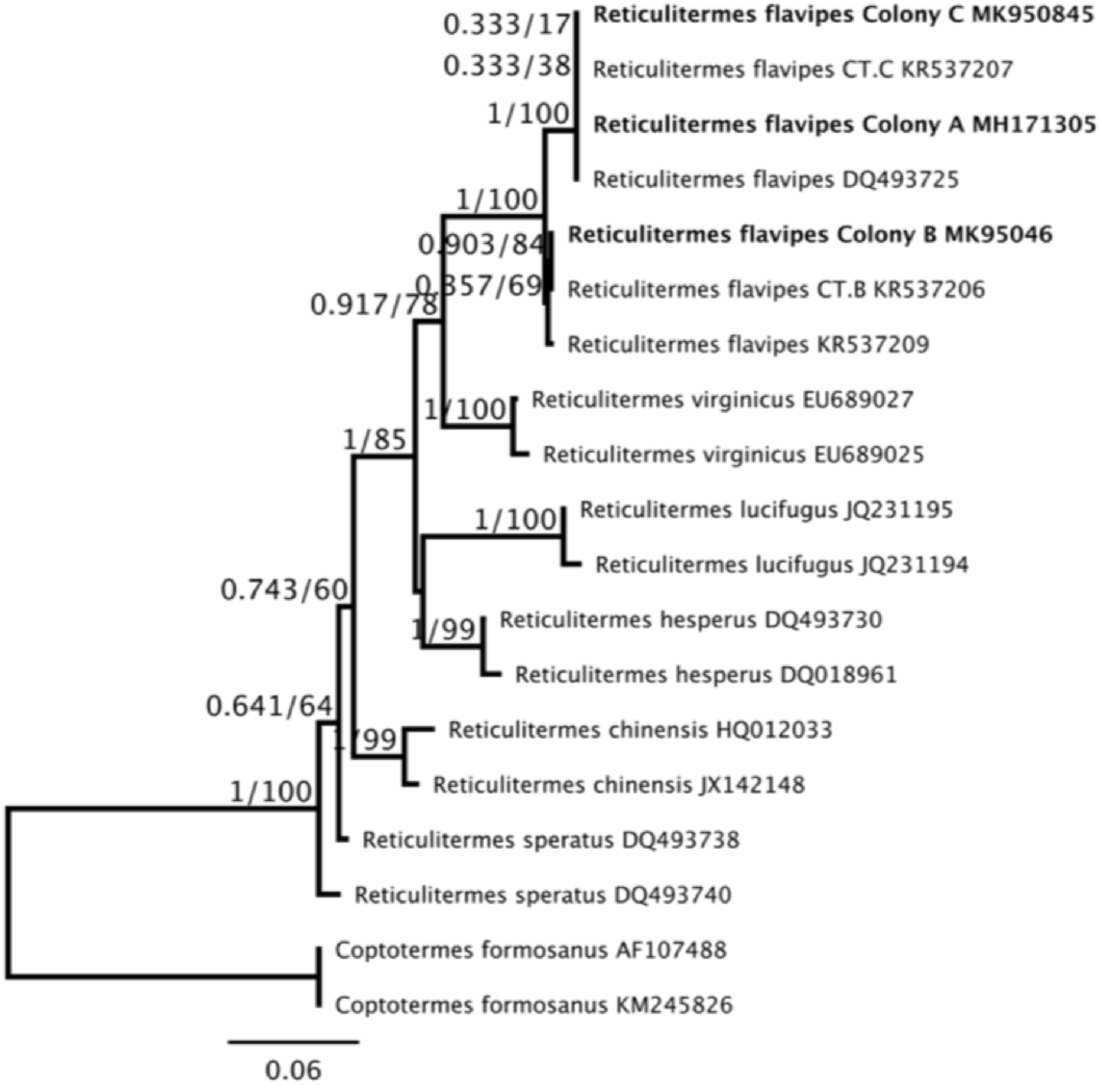
Phylogenetic tree of termite mitochondrial cytochrome oxidase II gene sequences. Sequences were aligned to references with MUSCLE and a Maximum likelihood (ML) phylogenetic tree was made using IQ-Tree with substitution model TIM2+G4. Sequences obtained from termites used in this study (Bold) clustered within the *R. flavipes* clade. Support values represent the Bayesian posterior probability and Bootstrap support values respectively.

**S4 Figure.**
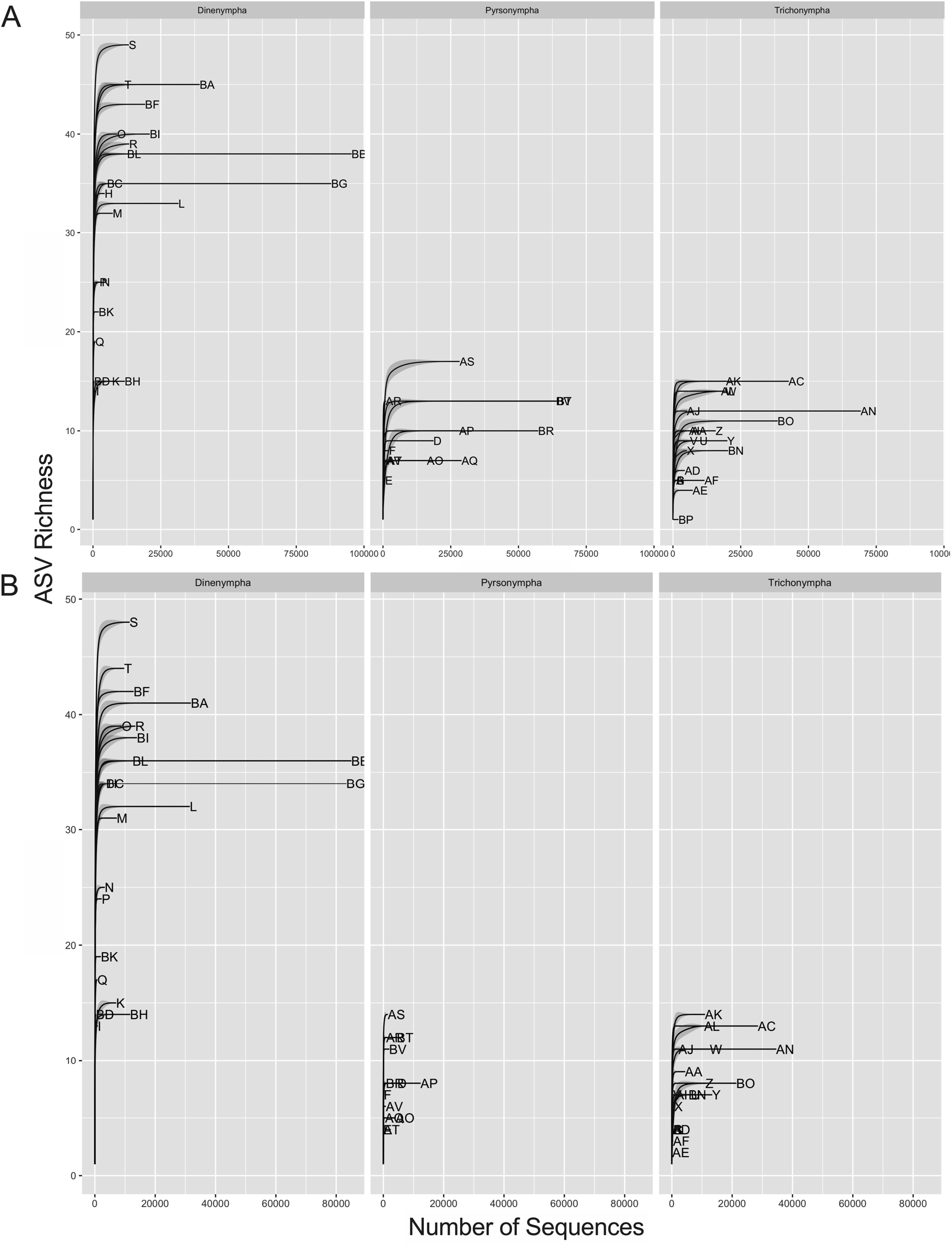
Rarefaction analysis of single-cell amplicon samples. Analysis was done using the “ggrare” function of ggplot2 samples with (A) and without (B) sequences of ‘*Candidatus* Endomicrobium’. Curve tips represent the cell’s identification.

**S5 Figure.**
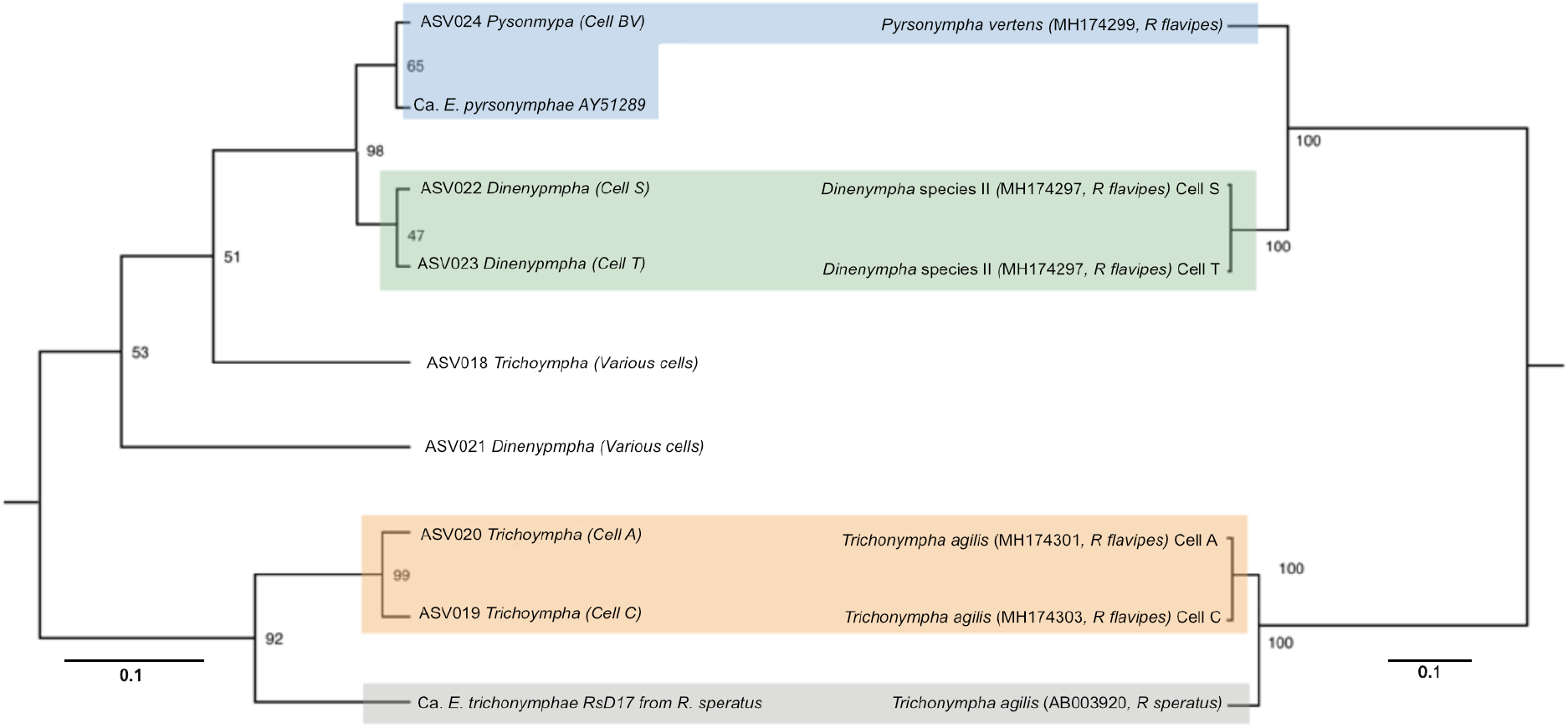
Congruent phylogenies between endosymbiotic ASVs and their protist hosts. *Endomicrobium* ASVs were aligned to the V4 regions of reference *Endomicrobium* species (left tree). The phylogenetic tree was made using Jukes-Cantor Distance Model with UPGMA as implemented in Geneious v9. The 18S rRNA genes from single protist hosts, and one reference sequence were aligned, and the tree was made using Jukes-Cantor Distance Model with UPGMA as implemented in Geneious v9 (right tree). Support values are Bootstrap percentage values. Sequences are color coded such endosymbionts are colored the same as their hosts. In the left tree, *Endomicrobium* ASVs are given host protist designations, and a host cell identification is given. These host cells contained the ASV as the majority, or only, member of the *Endomicrobium* associated with that cell. Other host cells (not indicated here but detailed in S1 File) also contained the *Endomicrobium* ASVs mapped here as sole, or majority, members of the association. Sequences of the *Endomicrobium* ASVs can be found in S1 File.

**S6 Figure.**
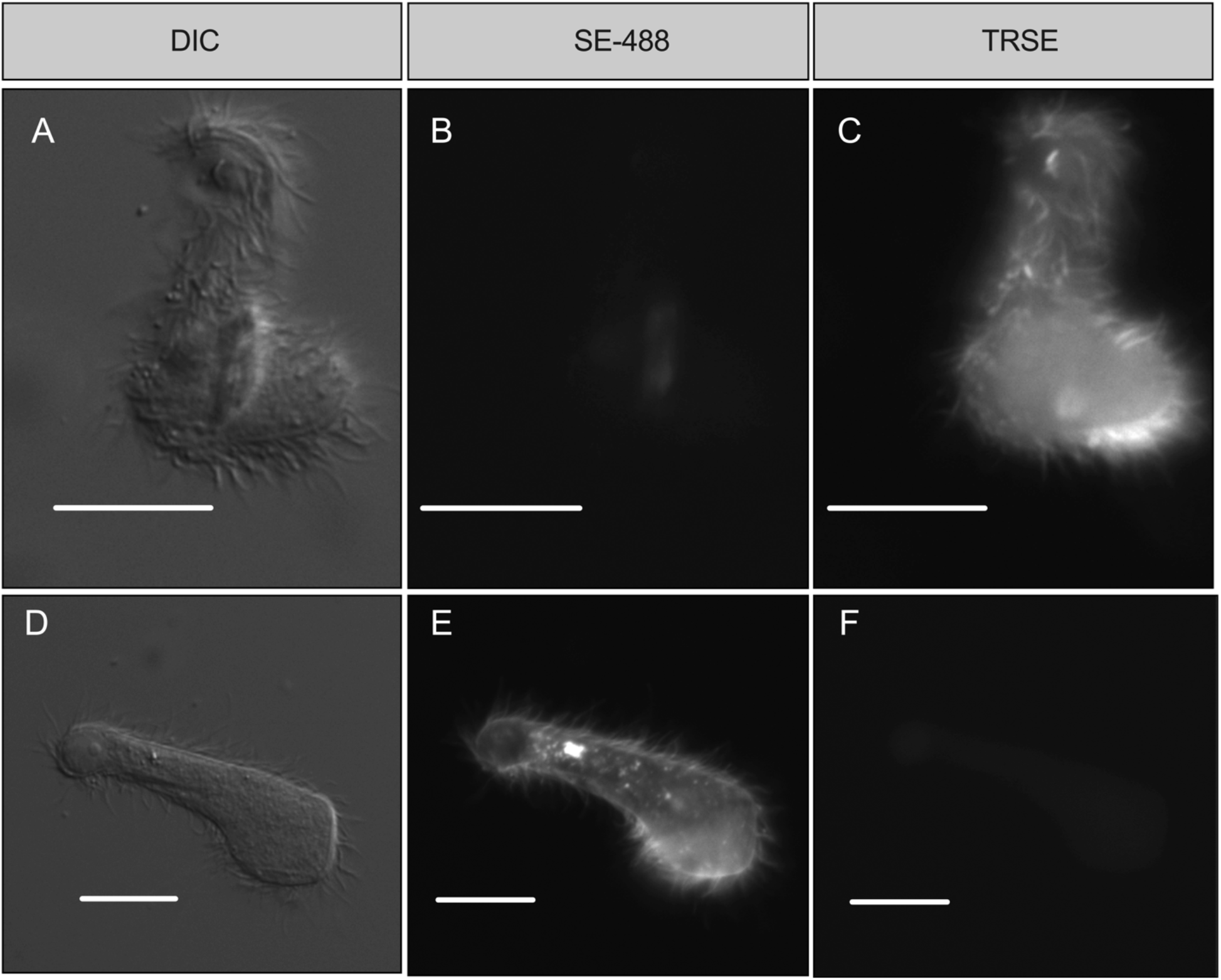
Non-mixed controls for fluorescent assays. Representative micrographs of protist cells from non-mixed control samples from fluorescent assays. Micrographs represent DIC (A and D) SE-488 fluorescence (B and E) and TRSE fluorescence (C and D). Control samples always maintained homogenous fluorescence. Micrographs were taken post 12 hours from the start of assays and scale bars represent 20μm.

## References

1. Brune A. Symbiotic digestion of lignocellulose in termite guts. Nat Rev Microbiol. 2014;12:168–80. Available from: http://www.ncbi.nlm.nih.gov/pubmed/24487819

2. Ohkuma M. Symbioses of flagellates and prokaryotes in the gut of lower termites. Trends Microbiol. 2008;16:345–52.

3. Noda S, Ohkuma M, Yamada A, Hongoh Y, Kudo T. Phylogenetic position and in situ identification of ectosymbiotic spirochetes on protists in the termite gut. Appl Environ Microbiol. 2003;69(1):625–33.

4. Iida T, Ohkuma M, Ohtoko K, Kudo T. Symbiotic spirochetes in the termite hindgut: Phylogenetic identification of ectosymbiotic spirochetes of oxymonad protists. FEMS Microbiol Ecol. 2000;34:17–26.

5. Hongoh Y, Sato T, Noda S, Ui S, Kudo T, Ohkuma M. *Candidatus* Symbiothrix dinenymphae: Bristle-like Bacteroidales ectosymbionts of termite gut protists. Environ Microbiol. 2007;9:2631–5.

6. Radek R, Tischendorf G. Bacterial adhesion to different termite flagellates: Ultrastructural and functional evidence for distinct molecular attachment modes. Protoplasma. 1999;207:43–53.

7. Sato T, Hongoh Y, Noda S, Hattori S, Ui S, Ohkuma M. *Candidatus* Desulfovibrio trichonymphae, a novel intracellular symbiont of the flagellate *Trichonympha agilis* in termite gut. Environ Microbiol. 2009;11(4):1007–15. Available from: http://www.ncbi.nlm.nih.gov/pubmed/19170725

8. Utami YD, Kuwahara H, Igai K, Murakami T, Sugaya K, Morikawa T, et al. Genome analyses of uncultured TG2/ZB3 bacteria in ‘Margulisbacteria’ specifically attached to ectosymbiotic spirochetes of protists in the termite gut. ISME J. 2019;13(2):455–67. Available from: http://dx.doi.org/10.1038/s41396-018-0297-4

9. Yuki M, Kuwahara H, Shintani M, Izawa K, Sato T, Starns D, et al. Dominant ectosymbiotic bacteria of cellulolytic protists in the termite gut also have the potential to digest lignocellulose. Environ Microbiol. 2015;17:4942–53.

10. Stingl U, Radek R, Yang H, Brune A. “Endomicrobia”: Cytoplasmic symbionts of termite gut protozoa form a separate phylum of prokaryotes. Appl Environ Microbiol. 2005;71(3):1473–9.

11. Strassert JFH, Mikaelyan A, Woyke T, Brune A. Genome analysis of ‘ *Candidatus* Ancillula trichonymphae’’, first representative of a deep-branching clade of Bifidobacteriales, strengthens evidence for convergent evolution in flagellate endosymbionts.’ Environ Microbiol Rep. 2016;8(5):865–73. Available from: http://doi.wiley.com/10.1111/1758-2229.12451

12. Sato T, Kuwahara H, Fujita K, Noda S, Kihara K, Yamada A, et al. Intranuclear verrucomicrobial symbionts and evidence of lateral gene transfer to the host protist in the termite gut. ISME J. 2014;8:1008–19. Available from: http://www.ncbi.nlm.nih.gov/pubmed/24335826

13. Hongoh Y, Sharma VK, Prakash T, Noda S, Toh H, Taylor TD, et al. Genome of an endosymbiont coupling N2 fixation to cellulolysis within protist cells in termite gut. Science. 2008;322(5904):1108–9. Available from: http://www.ncbi.nlm.nih.gov/pubmed/19008447

14. Hongoh Y, Sharma VK, Prakash T, Noda S, Taylor TD, Kudo T, et al. Complete genome of the uncultured Termite Group 1 bacteria in a single host protist cell. PNAS. 2008;105(14):5555–60.

15. Ikeda-Ohtsubo W, Brune A. Cospeciation of termite gut flagellates and their bacterial endosymbionts: Trichonympha species and “*Candidatus* Endomicrobium trichonymphae.” Mol Ecol. 2009;18(2):332–42.

16. Zheng H, Dietrich C, L. Thompson C, Meuser K, Brune A. Population Structure of Endomicrobia in Single Host Cells of Termite Gut Flagellates (*Trichonympha* spp.). Microbes Environ. 2015;30(1):92–8. Available from: https://www.jstage.jst.go.jp/article/jsme2/advpub/0/advpub_ME14169/_article

17. Benjamino J, Graf J. Characterization of the Core and Caste-Specific Microbiota in the Termite, *Reticulitermes flavipes*. Front Microbiol. 2016;7:171. Available from: http://www.pubmedcentral.nih.gov/articlerender.fcgi?artid=4756164&tool=pmcentrez&rendertype=abstract

18. Ye W, Lee CY, Scheffrahn RH, Aleong JM, Su NY, Bennett GW, et al. Phylogenetic relationships of nearctic *Reticulitermes* species (Isoptera: Rhinotermitidae) with particular reference to *Reticulitermes arenincola* Goellner. Mol Phylogenet Evol. 2004;30(3):815–22.

19. Lewis JL, Forschler BT. Protist communities from four castes and three species of *Reticulitermes* (Isoptera : Rhinotermitidae). Ann Entomol Soc Am. 2004;97(6):1242–51. Available from: isi:000225335000014

20. Lewis JL, Forschler BT. A nondichotomous key to protist species identification of *Reticulitermes* (Isoptera: Rhinotermitidae). Ann Entomol Soc Am. 2006;99(6):1028–33. Available from: http://aesa.oxfordjournals.org/content/99/6/1028.abstract

21. Maekawa K, Lo N, Kitade O, Miura T, Matsumoto T. Molecular phylogeny and geographic distribution of wood-feeding cockroaches in East Asian Islands. Mol Phylogenet Evol. 1999;13(2):360–76.

22. Trager W. The Cultivation of a Cellulose-Digesting Flagellate, *Trichomonas Termopsidis*, and of Certain other Termite Protozoa. Biol Bull. 1934;66(2):182–90.

23. Ohkuma M, Ohtoko K, Grunau C, Moriya S, Kudo T. Phylogenetic identification of the symbiotic hypermastigote *Trichonympha agilis* in the hindgut of the termite *Reticulitermes speratus* based on small-subunit rRNA sequence. J Eukaryot Microbiol. 1998;45(4):439–44.

24. Caporaso JG, Lauber CL, Walters WA, Berg-Lyons D, Huntley J, Fierer N, et al. Ultra-high-throughput microbial community analysis on the Illumina HiSeq and MiSeq platforms. ISME J. 2012;6(8):1621–4. Available from: http://dx.doi.org/10.1038/ismej.2012.8

25. Tikhonenkov D V., Janouškovec J, Keeling PJ, Mylnikov AP. The Morphology, Ultrastructure and SSU rRNA Gene Sequence of a New Freshwater Flagellate, *Neobodo borokensis* n. sp. (Kinetoplastea, Excavata). J Eukaryot Microbiol. 2016;63(2):220–32.

26. Callahan BJ, McMurdie PJ, Rosen MJ, Han AW, Johnson AJA, Holmes SP. DADA2: High-resolution sample inference from Illumina amplicon data. Nat Methods. 2016;13(7):581–3.

27. Callahan BJ, McMurdie PJ, Holmes SP. Exact sequence variants should replace operational taxonomic units in marker-gene data analysis. ISME J. 2017;11(12):2639–43. Available from: http://dx.doi.org/10.1038/ismej.2017.119

28. Quast C, Pruesse E, Yilmaz P, Gerken J, Schweer T, Yarza P, et al. The SILVA ribosomal RNA gene database project: Improved data processing and web-based tools. Nucleic Acids Res. 2013;41(D1):590–6.

29. Davis NM, Proctor DM, Holmes SP, Relman DA, Callahan BJ. Simple statistical identification and removal of contaminant sequences in marker-gene and metagenomics data. Microbiome. 2018;6(1):226. Available from: https://doi.org/10.1186/s40168-018-0605-2

30. McMurdie PJ, Holmes S. Phyloseq: An R Package for Reproducible Interactive Analysis and Graphics of Microbiome Census Data. PLoS One. 2013;8(4).

31. de Goffau MC, Lager S, Salter SJ, Wagner J, Kronbichler A, Charnock-Jones DS, et al. Recognizing the reagent microbiome. Nat Microbiol. 2018;3(8):851–3. Available from: http://dx.doi.org/10.1038/s41564-018-0202-y

32. Oksanen AJ, Kindt R, Legendre P, Hara BO, Simpson GL, Stevens MHH, et al. The vegan Package; Community Ecology Package (Version 1.15-1). 2008;

33. Wickham H. Ggplot2. Wiley Interdiscip Rev Comput Stat. 2011;3(2):180–5.

34. Edgar RC. MUSCLE: Multiple sequence alignment with high accuracy and high throughput. Nucleic Acids Res. 2004;32(5):1792–7.

35. Nguyen LT, Schmidt HA, Von Haeseler A, Minh BQ. IQ-TREE: A fast and effective stochastic algorithm for estimating maximum-likelihood phylogenies. Mol Biol Evol. 2015;32(1):268–74.

36. Pedro M a De, Grünfelder CG, Gru CG. Restricted Mobility of Cell Surface Proteins in the Polar Regions of *Escherichia coli*. J. Bacteriol. 2004;186(9).

37. Ohkuma M, Sato T, Noda S, Ui S, Kudo T, Hongoh Y. The candidate phylum “Termite Group 1” of bacteria: Phylogenetic diversity, distribution, and endosymbiont members of various gut flagellated protists. FEMS Microbiol Ecol. 2007;60(3):467–76.

38. McFall-Ngai MJ. The Importance of Microbes in Animal Development: Lessons from the Squid-Vibrio Symbiosis. Annu Rev Microbiol. 2014; Available from: http://www.ncbi.nlm.nih.gov/pubmed/24995875

39. Bright M, Bulgheresi S. A complex journey: transmission of microbial symbionts. Nat Rev Microbiol. 2010;8(3):218–30. Available from: http://dx.doi.org/10.1038/nrmicro2262%5Cnpapers3://publication/doi/10.1038/nrmicro2262

40. Peterson BF, Stewart HL, Scharf ME. Quantification of symbiotic contributions to lower termite lignocellulose digestion using antimicrobial treatments. Insect Biochem Mol Biol. 2015;59:80–8. Available from: http://dx.doi.org/10.1016/j.ibmb.2015.02.009

41. Kodama Y, Fujishima M. Cycloheximide Induces Synchronous Swelling of Perialgal Vacuoles Enclosing Symbiotic *Chlorella vulgaris* and Digestion of the Algae in the Ciliate *Paramecium bursaria*. Protist. 2008;159(3):483–94.

42. Corno G, Jürgens K. Direct and indirect effects of protist predation on population size structure of a bacterial strain with high phenotypic plasticity. Appl Environ Microbiol. 2006;72(1):78–86. Available from: http://aem.asm.org/content/72/1/78.full.pdf

43. Boulos L, Prévost M, Barbeau B, Coallier J, Desjardins R. LIVE/DEAD(®) BacLight(TM): Application of a new rapid staining method for direct enumeration of viable and total bacteria in drinking water. J Microbiol Methods. 1999;37(1):77–86.

44. Waidele L, Korb J, Voolstra CR, Künzel S, Dedeine F, Staubach F. Differential ecological specificity of protist and bacterial microbiomes across a set of termite species. Front Microbiol. 2017;8:1–13.

45. James ER, Tai V, Scheffrahn RH, Keeling PJ. Trichonympha burlesquei n. sp. From *Reticulitermes virginicus* and evidence against a cosmopolitan distribution of *Trichonympha agilis* in many termite hosts. Int J Syst Evol Microbiol. 2013;63(PART10):3873–6.

46. Bourguignon T, Lo N, Dietrich C, Šobotník J, Sidek S, Roisin Y, et al. Rampant Host Switching Shaped the Termite Gut Microbiome. Curr Biol. 2018;28(4):649–654.e2.

47. Treitli SC, Kolisko M, Husník F, Keeling PJ, Hampl V. Revealing the metabolic capacity of *Streblomastix strix* and its bacterial symbionts using singlecell metagenomics. Proc Natl Acad Sci U S A. 2019;116(39):19675–84.

48. Graber JR, Breznak J a. Physiology and Nutrition of *Treponema primitia*, an H2 / CO2 -Acetogenic Spirochete from Termite Hindguts. Appl Environ Microbiol. 2004;70(3):1307–14.

49. Yamin M a., Trager W. Cellulolytic Activity of an Axenically-cultivated Termite Flagellate, *Trichomitopsis termopsidis*. J Gen Microbiol. 1979;113:417–20.

50. Odelson D a., Breznak J a. Nutrition and growth characteristics of *Trichomitopsis termopsidis*, a cellulolytic protozoan from termites. Appl Environ Microbiol. 1985;49(3):614–21.

51. Tai V, Carpenter KJ, Weber PK, Nalepa CA, Perlman SJ, Keeling PJ. Genome evolution and nitrogen-fixation in bacterial ectosymbionts of a protist inhabiting wood-feeding cockroaches. Appl Environ Microbiol. 2016;82(15):4682–95. Available from: http://aem.asm.org/lookup/doi/10.1128/AEM.00611-16

52. Jasso-Selles DE, De Martini F, Freeman KD, Garcia MD, Merrell TL, Scheffrahn RH, et al. The parabasalid symbiont community of *Heterotermes aureus*: Molecular and morphological characterization of four new species and reestablishment of the genus *Cononympha*. Eur J Protistol. 2017;61:48–63. Available from: http://dx.doi.org/10.1016/j.ejop.2017.09.001

