## Supplementary material for "Single-Cell Amplicon Sequencing Reveals Community Structures and Transmission Trends of Protist-Associated Bacteria in a Termite Host": R script to repeat data analyses

# This R Script was pieced together by: Michael E. Stephens, University of Connecicut, Storrs CT, USA
# Used to analyze bacterial ASVs associated with protists isolated from the hindgut of the termite, Reticulitermes flavipes
# This script is provided for the reproducibility of all ASV-related analyses used in this study
# 
# This Pipeline uses raw R1 and R2 read files (.fastq format) for 16S rRNA gene (V4 region) amplicons to generate ASVs, decontaminate (if controls were used), filter, and generate graphs
# dada2 pipiline taken follows tutorial: https://benjjneb.github.io/dada2/tutorial.html
# Other tutorials and refernces are provided throughout this R script, they are all publically avialiable 

# USER MUST SET PATHS TO APPROPIATE DIRECTORIES FOR READS, TAXONOMY FILES, MAPPING FILE, AND ALL OUTPUT FILES

# Step 1 #####################################################################################
# Load all packages #########################################################################

library(dada2); packageVersion("dada2")
library(phyloseq); packageVersion("phyloseq")
library(ggplot2); packageVersion("ggplot2")
library(vegan)
library(decontam)

# Packages for some downstream analyses, following: https://www.bioconductor.org/help/course-materials/2017/BioC2017/Day1/Workshops/Microbiome/MicrobiomeWorkflowII.html

library("knitr")
library("BiocStyle")
.cran_packages <- c("ggplot2", "gridExtra")
.bioc_packages <- c("dada2", "phyloseq", "DECIPHER", "phangorn")
.inst <- .cran_packages %in% installed.packages()
if(any(!.inst)) {
  install.packages(.cran_packages[!.inst])
}
.inst <- .bioc_packages %in% installed.packages()
if(any(!.inst)) {
  source("http://bioconductor.org/biocLite.R")
  biocLite(.bioc_packages[!.inst], ask = F)
}
sapply(c(.cran_packages, .bioc_packages), require, character.only = TRUE)

# Step 2 ####################################################################################
#Load in Read files and visualize quality ##################################################################################################

# Set Path to direcotry with un-zipped read files (.fastq)

path <-""
list.files(path)

# Forward and reverse fastq filenames have format: SAMPLENAME_R1_001.fastq and SAMPLENAME_R2_001.fastq

fnFs <- sort(list.files(path, pattern="_R1_001.fastq", full.names = TRUE))
fnRs <- sort(list.files(path, pattern="_R2_001.fastq", full.names = TRUE))

# Extract sample names, assuming filenames have format: SAMPLENAME_XXX.fastq

sample.names <- sapply(strsplit(basename(fnFs), "_"), `[`, 1)

# Visulaize quality profiles of Forward Reads

plotQualityProfile(fnFs[1:82]) # Set to the number of reads you want to see

## Scale for 'y' is already present. Adding another scale for 'y', which will replace the existing scale.

# Visulaize quality profiles of Reverse Reads

plotQualityProfile(fnRs[1:82]) # Set to the number of reads you want to see


#Step 3 #########################################################################################
#Filter reads based on size, Trim, learn error rates, Dereplicate, and Merge paired Reads ####################################################

# Place filtered files in filtered/ subdirectory

filtFs <- file.path(path, "filtered", paste0(sample.names, "_F_filt.fastq.gz"))
filtRs <- file.path(path, "filtered", paste0(sample.names, "_R_filt.fastq.gz"))

# Filter and Trim

out <- filterAndTrim(fnFs, filtFs, fnRs, filtRs, truncLen=c(240,160),
                     maxN=0, maxEE=c(2,2), truncQ=2, rm.phix=TRUE,
                     compress=TRUE, multithread=TRUE) # On Windows set multithread=FALSE
head(out)

# Learn Forward and Reverse Error Rates

errF <- learnErrors(filtFs, multithread=TRUE)
errR <- learnErrors(filtRs, multithread=TRUE)

# Visualize Forward Error Rates

plotErrors(errF, nominalQ=TRUE)+
  ggsave("forward_errors.png", path = "")

plotErrors(errR, nominalQ =TRUE)+
  ggsave("reverse_errors.png", path = "")

# Derepilication
# Dereplication combines all identical sequencing reads into into “unique sequences” with a corresponding “abundance” equal to the number of reads with that unique sequence. 
# Dereplication substantially reduces computation time by eliminating redundant comparisons.

derepFs <- derepFastq(filtFs, verbose=TRUE)
derepRs <- derepFastq(filtRs, verbose=TRUE)

# Name the derep-class objects by the sample names

names(derepFs) <- sample.names
names(derepRs) <- sample.names

# Apply the core sample inference algorithm (dada) to the dereplicated data

dadaFs <- dada(derepFs, err=errF, multithread=TRUE)
dadaRs <- dada(derepRs, err=errR, multithread=TRUE)

# Merge Paired Reads
mergers <- mergePairs(dadaFs, derepFs, dadaRs, derepRs, verbose=TRUE)

# Inspect the merger data.frame from the first sample
head(mergers[[1]])

# Construct Sequence Table
seqtab <- makeSequenceTable(mergers)
dim(seqtab)
seqtab2 <- seqtab[,nchar(colnames(seqtab)) %in% seq(250,256)]

# Inspect distribution of sequence lengths
table(nchar(getSequences(seqtab)))
table(nchar(getSequences(seqtab2)))


# Step 4 ####################################################################################
# Remove Chimeras ############################################################################################

# Remove Chimeras
seqtab.nochim <- removeBimeraDenovo(seqtab2, method="consensus", multithread=TRUE, verbose=TRUE)
dim(seqtab.nochim)

# % of Reads Remaining 
sum(seqtab.nochim)/sum(seqtab2)

# Track Reads Through the Pipleline so far
getN <- function(x) sum(getUniques(x))
track <- cbind(out, sapply(dadaFs, getN), sapply(dadaRs, getN), sapply(mergers, getN), rowSums(seqtab.nochim))

# If processing a single sample, remove the sapply calls: e.g. replace sapply(dadaFs, getN) with getN(dadaFs)
colnames(track) <- c("input", "filtered", "denoisedF", "denoisedR", "merged", "nonchim")
rownames(track) <- sample.names
(track)

# Step 5 ##########################################################################################
# Assign Taxonomy, generate initial ASV counts and taxonomy tables ###########################################################################################

# Assign Taxonomy

taxa <- assignTaxonomy(seqtab.nochim, "silva_nr_v132_train_set.fa", multithread=TRUE) #Path to taxonomy training set
taxa <- addSpecies(taxa,"silva_species_assignment_v132.fa") #Path to second (species) taxonomy training set

# Inspect Taxonomy

taxa.print <- taxa # Removing sequence rownames for display only
rownames(taxa.print) <- NULL
head(taxa.print)

# Step 6 ##########################################################################################
# Align seqs, make a phylogenetic tree, and combine information into a phyloseq object #########################################################

# Follows tutoiral found here: https://www.bioconductor.org/help/course-materials/2017/BioC2017/Day1/Workshops/Microbiome/MicrobiomeWorkflowII.html
# This can take a few minutes..or hours...

seqs <- getSequences(seqtab.nochim)
names(seqs) <- seqs # This propagates to the tip labels of the tree
alignment <- AlignSeqs(DNAStringSet(seqs), anchor=NA,verbose=FALSE)

phangAlign <- phyDat(as(alignment, "matrix"), type="DNA")
dm <- dist.ml(phangAlign)
treeNJ <- NJ(dm) # Note, tip order != sequence order
fit = pml(treeNJ, data=phangAlign)
fitGTR <- update(fit, k=4, inv=0.2)
fitGTR <- optim.pml(fitGTR, model="GTR", optInv=TRUE, optGamma=TRUE,
                    rearrangement = "stochastic", control = pml.control(trace = 0))
detach("package:phangorn", unload=TRUE)

# Combine data into a Phyloseq Object
# Fix or confirm row names are sample name

samdf <- read.csv("")  # samdf = mapping file (.csv)
all(rownames(seqtab.nochim) %in% samdf$SampleID) # TRUE
rownames(samdf) <- samdf$SampleID
keep.cols <- c("sample_or_control", "Host_species", "Host_genus", "Host_phyla", "cell_id", "sample_collection_date", "termite_id", "termite_colony", "lab_colony_id", "colony_collection_date")
samdf <- samdf[rownames(seqtab.nochim), keep.cols]

#Make Phyloseq Object (ps)

ps <- phyloseq(otu_table(seqtab.nochim, taxa_are_rows=FALSE), 
               sample_data(samdf), 
               tax_table(taxa),phy_tree(fitGTR$tree))
ps

# Inspect ASV table #

# Extract abundance matrix from the unfiltered phyloseq object
OTU1 = as(otu_table(ps), "matrix")
# transpose if necessary
if(taxa_are_rows(ps)){OTU1 <- t(OTU1)}
# Coerce to data.frame
OTUdf = as.data.frame(OTU1)
write.table(OTUdf, file = "", sep = " ")


# Step 7 ##########################################################################################
# Decontaminate samples using package "decontam" and method "prevalence", must have negative control samples ###########################################
# Uses a chi-square statistic on the 2x2 contingency table of presence/absence in true samples and in negative controls
# Fisher’s exact test may be used instead of chi-square statistic if there are too few samples
# Method descriped in :Davis NM, Proctor DM, Holmes SP, Relman DA, Callahan BJ. Simple statistical identification and removal of contaminant sequences in marker-gene and metagenomics data. Microbiome. 2018;6: 226. doi:10.1186/s40168-018-0605-2

# Following tutorital from: https://benjjneb.github.io/decontam/vignettes/decontam_intro.html

# Inspect library size

df <- as.data.frame(sample_data(ps)) # Put sample_data into a ggplot-friendly data.frame
df$LibrarySize <- sample_sums(ps)
df <- df[order(df$LibrarySize),]
df$Index <- seq(nrow(df))
ggplot(data=df, aes(x=Index, y=LibrarySize, color=sample_or_control)) + geom_point()+
  ggsave("library_size.png", path = "")

# Indentify contaminat using the prevalence method

sample_data(ps)$is.neg <- sample_data(ps)$sample_or_control == "control"
contamdf.prev <- isContaminant(ps, method="prevalence", neg= "is.neg")
table(contamdf.prev$contaminant)

head(which(contamdf.prev$contaminant))

contamdf.prev04 <- isContaminant(ps, method="prevalence", neg="is.neg", threshold=0.35) 
table(contamdf.prev04$contaminant)

# Make phyloseq object of presence-absence in negative controls

ps.neg <- prune_samples(sample_data(ps)$sample_or_control == "control", ps)
ps.neg.presence <- transform_sample_counts(ps.neg, function(abund) 1*(abund>0))

# Make phyloseq object of presence-absence in true positive samples

ps.pos <- prune_samples(sample_data(ps)$sample_or_control == "sample", ps)
ps.pos.presence <- transform_sample_counts(ps.pos, function(abund) 1*(abund>0))

# Make data.frame of prevalence in positive and negative samples

df.pres <- data.frame(prevalence.pos=taxa_sums(ps.pos.presence), prevalence.neg=taxa_sums(ps.neg.presence),
                      Contaminate=contamdf.prev$contaminant)
ggplot(data=df.pres,aes(x=prevalence.neg, y=prevalence.pos, color=Contaminate)) + geom_point()+
  xlab("Prevalence (Negative Controls)") + ylab("Prevalence (Protist Samples)")+
  ggsave("contam_prev.png", path = "")

# Remove contaminants

ps_noncontam <- prune_taxa(!contamdf.prev04$contaminant, ps)
ps_noncontam


# Remove control samples from phyloseq object

ps_nocontrols <- prune_samples(sample_data(ps_noncontam)$sample_or_control == "sample", ps_noncontam)
ps_nocontrols


# Step 8 ##################################################################################################################
# Perform Taxonomic filtering (i.e what do you expect base on a-priori knowledge of your samples; Ecological plausibility) ##################################
# Following contamination identification guidelines published in: de Goffau MC, Lager S, Salter SJ, Wagner J, Kronbichler A, Charnock-Jones DS, et al. Recognizing the reagent microbiome. Nat Microbiol. Springer US; 2018;3: 851–853. doi:10.1038/s41564-018-0202-y
# Create table, number of features for each phyla

table(tax_table(ps_nocontrols)[, "Phylum"], exclude = NULL)

# Remove uncharacterized seqs
ps_nocontrols_b <- subset_taxa(ps_nocontrols, !is.na(Phylum) & !Phylum %in% c("", "uncharacterized"))
ps_nocontrols_b

# Compute prevalence of each feature, store as data.frame
prevdf = apply(X = otu_table(ps_nocontrols_b),
               MARGIN = ifelse(taxa_are_rows(ps_nocontrols_b), yes = 1, no = 2),
               FUN = function(x){sum(x > 0)})

# Add taxonomy and total read counts to this data.frame
prevdf = data.frame(Prevalence = prevdf,
                    TotalAbundance = taxa_sums(ps_nocontrols_b),
                    tax_table(ps_nocontrols_b))

#Are there phyla that are comprised of mostly low-prevalence features? Compute the total and average prevalences of the features in each phylum
plyr::ddply(prevdf, "Phylum", function(df1){cbind(mean(df1$Prevalence),sum(df1$Prevalence))})

# Define phyla to filter
filterPhyla = c("Acidobacteria", "Actinobacteria", "Cyanobacteria", "Euryarchaeota", "Firmicutes", "Fusobacteria", "Gemmatimonadetes", "Planctomycetes", "Tenericutes", "Chloroflexi", "Chlamydiae")

# Filter entries with Phyla list from above
ps1 = subset_taxa(ps_nocontrols_b, !Phylum %in% filterPhyla)
ps1

# Define Orders to filter
filterOrders = c("Rhizobiales", "Betaproteobacteriales", "Pseudomonadales", "Sphingomonadales", "Xanthomonadales", "Rickettsiales", "Rhodobacterales", "Rhodospirillales", "Sphingobacteriales", "Enterobacteriales", "Salinisphaerales", "Caedibacterales", "Myxococcales", "Kordiimonadales", "Xanthomonadales", "Flavobacteriales", "Chitinophagales", "Cytophagales", "Caulobacterales", "Holosporales", "Oceanospirillales","Oligoflexales", "Paracaedibacterales")

# Filter entries with Order list from above
ps1a = subset_taxa(ps1, !Order %in% filterOrders)
ps1a

# Define Genera to filter
filterGenera = c("Aquabacterium", "Ralstonia", "Sphingomonas", "Pelomonas", "Escherichia/Shigella", "Bradyrhizobium", "Methylobacterium", "Stenotrophomonas", "Caulobacter", "Allorhizobium-Neorhizobium-Pararhizobium-Rhizobium
", "Pelomonas", "Cupriavidus", "Brucella", "Klebsiella", "Curvibacter", "Vanovorax", "Roseomonas", "Acidibacter", "Vibrionimonas", "Ruegeria")

# Filter entries with Genera list from above
ps1b = subset_taxa(ps1a, !Genus %in% filterGenera)
ps1b

# Step 9 #################################################################################################
# Filter based on prevalence in protist samples, i.e remove ASVs in fewer than X% of samples
# Define prevalence threshold as X% (i.e 5%, 0.05) of total samples
                 
prevalenceThreshold = 0.05263158 * nsamples(ps1b)  # threshold x # samples = minimum of 3 protist samples
prevalenceThreshold


# Subset to the remaining phyla
prevdf1 = subset(prevdf, Phylum %in% get_taxa_unique(ps1b, "Phylum"))
ggplot(prevdf1, aes(TotalAbundance, Prevalence / nsamples(ps1b),color=Phylum)) +
  # Include a guess for parameter
  geom_hline(yintercept = 0.1, alpha = 0.5, linetype = 2) +  geom_point(size = 2, alpha = 0.7) +
  scale_x_log10() +  xlab("Total Abundance") + ylab("Prevalence [Frac. Samples]") +
  facet_wrap(~Phylum) + theme(legend.position="none") +
  ggsave("tax_prevalence_nonfiltered.png", path = "")

# Execute prevalence filter, using `prune_taxa()` function

keepTaxa = rownames(prevdf1)[(prevdf1$Prevalence >= prevalenceThreshold)]
ps2 = prune_taxa(keepTaxa, ps1b)
ps2

  
# How many genera would be present after prevalence filtering?

length(get_taxa_unique(ps2, taxonomic.rank = "Genus"))

# Subset to the remaining phyla after prevalence filtering

prevdf2 = subset(prevdf, Phylum %in% get_taxa_unique(ps2, "Phylum"))
ggplot(prevdf2, aes(TotalAbundance, Prevalence / nsamples(ps2),color=Phylum)) +
  # Include a guess for parameter
  geom_hline(yintercept = 0.1, alpha = 0.5, linetype = 2) +  geom_point(size = 2, alpha = 0.7) +
  scale_x_log10() +  xlab("Total Abundance") + ylab("Prevalence [Frac. Samples]") +
  facet_wrap(~Phylum) + theme(legend.position="none") +
  ggsave("tax_prevalence_filtered.png", path = "")
  
# Step 10 ##############################################################################################

# Transform to relative abundance. Save as new object.

ps2ra = transform_sample_counts(ps2, function(x){(x / sum(x))*100})


  
#Step 11 ###################################################################################
# Remove samples with fewer than X numbers of reads left at this stage of filtering, remember these are low biomass samples that have been filtered prior to this step

ps2_low_samples_removed = prune_samples(sample_sums(ps2)>=500, ps2) 
ps2_low_samples_removed


#Step 12 ######################################################################################
# Filter ASVs that are below a minium relative abundance threshold per sample
                                                                                                     
ps_prunedSet = prune_taxa((taxa_sums(ps2_low_samples_removed)/sum(taxa_sums(ps2_low_samples_removed))) >=1e-5, ps2_low_samples_removed)
ps_prunedSet

#inspect Endomicrobium
endo <- subset_taxa(ps_prunedSet, Genus=="Candidatus_Endomicrobium")
endo

# Extract abundance matrix from the phyloseq object
OTU_endo = as(otu_table(endo), "matrix")
# transpose if necessary
if(taxa_are_rows(endo)){OTU_endo <- t(OTU_endo)}
# Coerce to data.frame
OTU_endo_df = as.data.frame(OTU_endo)
write.table(OTU_endo_df, file = "", sep = " ")

# Transform counts 

ps_prunedSet_ra = transform_sample_counts(ps_prunedSet, function(x){(x / sum(x))*100})


# Step 13 ###################################################################################
# Ordination of remaining samples and thier ASVs #############################################

asv_nmds3 <- ordinate(
  physeq = ps_prunedSet_ra, 
  method = "NMDS",
  distance = "bray" # Able to sub in different methods here for distace ie. "bray" "wunifrac" "unifrac"
)
nmds_plot_pruned = plot_ordination(
  physeq = ps_prunedSet_ra,
  ordination = asv_nmds3,
  label = "cell_id", 
  color = "Host_genus",
  shape = "termite_colony",
  title = "Ordination of Bacterial Communities of Hindgut Protists (Bray-Curtis)"
)+
  stat_ellipse(type = "t", linetype = 5) +                         #type = "t" = multivariate t distribution, can also be "norm"
  labs(col = "Protist Genera", shape = "Termite Colonies")+
  ggsave("protists_nmds_bray_pruned.png", path = "" ) 
print(nmds_plot_pruned)

# Step 14 ######################################################################################################
# Make heatmap for samples and thier ASVs, separate by Protist Genus

# Make heatmap of pruned set
                                 #ps_prunedSet_B or ps_prunedSet_ra
heatmap_prunedSet = plot_heatmap(ps_prunedSet_ra, "NMDS", "bray", "cell_id", "Phylum", na.value = "grey90", low="#000033", high="#66CCFF")+
  geom_tile(colour="white",size=0.40)+
  #set a base size for all fonts
  theme_grey(base_size=8)+
  #theme options
  theme(
    #bold font for both axis text
    axis.text=element_text(face="bold"),
    #set thickness of axis ticks
    axis.ticks=element_line(size=0.4),
    #remove plot background
    plot.background=element_blank(),
    #remove plot border
    panel.border=element_blank())+
  facet_grid(~Host_genus, scale="free")+
  theme(strip.text.x = element_text(size = 12))+
theme(axis.text.x = element_text(size = 7, angle = 45))+
theme(legend.position = "bottom")
heatmap_prunedSet$scales$scales[[1]]$name <- "Protist Cells"
heatmap_prunedSet$scales$scales[[2]]$name <- "Bacterial ASVs (Phylum)"
print(heatmap_prunedSet)+
  ggsave("heatmap_prunedSet_ra.png", path = "" )


# Step 15 #############################################################################################
# Estimate and Plot Richness/ Alpha diversity #######

ps_prunedSet_alpha_est <- estimate_richness(ps_prunedSet, measures = c("Shannon", "Chao1", "Observed"))
write.csv(ps_prunedSet_alpha_est, file ="" )

alpha_p <- plot_richness(ps_prunedSet, x="Host_genus", measures=c("Observed", "Shannon", "Chao1"), color="termite_colony")+
  labs(col = "Termite Colonies", x = "Protist Genera")+
  ggsave("protist_richness_plots.png", path = "" )
(alpha_box <- alpha_p + geom_boxplot(data=alpha_p$data, aes(x = Host_genus, y=value, color=NULL), alpha=0.1))+
  ggsave("protist_richness_boxplots.png", path = "" )


#Step 16 #############################################################################################
# Arrange all final graphs (Heatmap, NMDS, Abundance) together
# Make figure

g_heatmap_plots_prunedSet <-arrangeGrob(heatmap_prunedSet, alpha_box, nmds_plot_pruned,                                    
                                         ncol = 3, nrow = 2, 
                                         layout_matrix = rbind(c(1,2), c(1,3), c(1,4)))
ggsave("heatmap_plots_prunedSetB.png", g_heatmap_plots_prunedSet, path = "")

# Step 19 #################################################################################################
# Run Adonis statistic on samples (Is there a difference between the protist Genera?)
# Extract Abundance matrix from phyloseq object

ASV_table_pruned = OTU1 = as(otu_table(ps_prunedSet_ra), "matrix")

# Transpose ASV table if needed

if(taxa_are_rows(ps_prunedSet_ra)){ASV_table_pruned <- t(ASV_table_pruned)}

# Coerce to data.fram

ASV_table_pruned_df = as.data.frame(ASV_table_pruned)
write.table(ASV_table_pruned_df, "", sep="\t", quote=F)


# Run adonis on ASV dataset
ASV_dist = phyloseq::distance(ps_prunedSet_ra, "bray")
ps_prunedSet_NMDS = ordinate(ps_prunedSet_ra, "NMDS", ASV_dist)
adonis(ASV_dist ~ Host_genus, as(sample_data(ps_prunedSet_ra), "data.frame"))

adonis(ASV_dist ~ termite_colony, as(sample_data(ps_prunedSet_ra), "data.frame"))

ps_trichonympha = subset_samples(ps_prunedSet_ra, Host_genus== "Trichonympha")
ps_trichonympha

ASV_dist = phyloseq::distance(ps_trichonympha, "bray")
ps_prunedSet_NMDS = ordinate(ps_trichonympha, "NMDS", ASV_dist)
adonis(ASV_dist ~ termite_colony, as(sample_data(ps_trichonympha), "data.frame"))

ps_pyrsonympha = subset_samples(ps_prunedSet_ra, Host_genus== "Pyrsonympha")
ps_pyrsonympha

ASV_dist = phyloseq::distance(ps_pyrsonympha, "bray")
ps_prunedSet_NMDS = ordinate(ps_pyrsonympha, "NMDS", ASV_dist)
adonis(ASV_dist ~ termite_colony, as(sample_data(ps_pyrsonympha), "data.frame"))

ps_dinenympha = subset_samples(ps_prunedSet_ra, Host_genus== "Dinenympha")
ps_dinenympha

ASV_dist = phyloseq::distance(ps_dinenympha, "bray")
ps_prunedSet_NMDS = ordinate(ps_dinenympha, "NMDS", ASV_dist)
adonis(ASV_dist ~ termite_colony, as(sample_data(ps_dinenympha), "data.frame"))


# Make bar charts

bar_plot = plot_bar(ps_prunedSet_ra, "cell_id", fill="Phylum")+ 
           facet_grid(~Host_genus, scale="free")
print(bar_plot)
ggsave("barplot_prunedSet.png", bar_plot, path = "")


#inspect Treponema ASVs
spiro_asvs <- subset_taxa(ps_prunedSet, Phylum=="Spirochaetes")
spiro_asvs

# Extract abundance matrix from the phyloseq object
OTU_spiro = as(otu_table(spiro_asvs), "matrix")
# transpose if necessary
if(taxa_are_rows(spiro_asvs)){OTU_spiro <- t(OTU_spiro)}
# Coerce to data.frame
OTU_spiro_df = as.data.frame(OTU_spiro)
write.table(OTU_spiro_df, file = "", sep = " ")


#inspect Candidatus_Symbiothrix ASVs
bacteroid_asvs <- subset_taxa(ps_prunedSet, Phylum=="Bacteroidetes")
bacteroid_asvs

# Extract abundance matrix from the phyloseq object
OTU_bacteroid = as(otu_table(bacteroid_asvs), "matrix")
# transpose if necessary
if(taxa_are_rows(bacteroid_asvs)){OTU_bacteroid <- t(OTU_bacteroid)}
# Coerce to data.frame
OTU_bacteroid_df = as.data.frame(OTU_bacteroid)
write.table(OTU_bacteroid_df, file = "", sep = " ")

#inspect Desulfovibrio ASVs
proteo_asvs <- subset_taxa(ps_prunedSet, Phylum=="Proteobacteria")
proteo_asvs

# Extract abundance matrix from the phyloseq object
OTU_proteo = as(otu_table(proteo_asvs), "matrix")
# transpose if necessary
if(taxa_are_rows(proteo_asvs)){OTU_proteo <- t(OTU_proteo)}
# Coerce to data.frame
OTU_proteo_df = as.data.frame(OTU_proteo)
write.table(OTU_proteo_df, file = "", sep = " ")

#inspect Verrucomicrobia ASVs
Verrucomicrobia_asvs <- subset_taxa(ps_prunedSet, Phylum=="Verrucomicrobia")
Verrucomicrobia_asvs

# Extract abundance matrix from the phyloseq object
OTU_Verrucomicrobia = as(otu_table(Verrucomicrobia_asvs), "matrix")
# transpose if necessary
if(taxa_are_rows(Verrucomicrobia_asvs)){OTU_Verrucomicrobia <- t(OTU_Verrucomicrobia)}
# Coerce to data.frame
OTU_Verrucomicrobia_df = as.data.frame(OTU_Verrucomicrobia)
write.table(OTU_Verrucomicrobia_df, file = "", sep = " ")

#inspect Margulisbacteria ASVs
Margulisbacteria_asvs <- subset_taxa(ps_prunedSet, Phylum=="Margulisbacteria")
Margulisbacteria_asvs

# Extract abundance matrix from the phyloseq object
OTU_Margulisbacteria = as(otu_table(Margulisbacteria_asvs), "matrix")
# transpose if necessary
if(taxa_are_rows(Margulisbacteria_asvs)){OTU_Margulisbacteria <- t(OTU_Margulisbacteria)}
# Coerce to data.frame
OTU_Margulisbacteria_df = as.data.frame(OTU_Margulisbacteria)
write.table(OTU_Margulisbacteria_df, file = "", sep = " ")

# Write taxonomy file
tax_prundedset <- tax_table(ps_prunedSet)

write.csv(tax_prundedset, file = "", sep = " " )

# inspect Synergistetes ASVs
Synergistetes_asvs <- subset_taxa(ps_prunedSet, Phylum=="Synergistetes")
Synergistetes_asvs

# Extract abundance matrix from the phyloseq object
OTU_Synergistetes = as(otu_table(Synergistetes_asvs), "matrix")
# transpose if necessary
if(taxa_are_rows(Synergistetes_asvs)){OTU_Synergistetes <- t(OTU_Synergistetes)}
# Coerce to data.frame
OTU_Synergistetes_df = as.data.frame(OTU_Synergistetes)
write.table(OTU_Synergistetes_df, file = "", sep = " ")

#inspect Endomicrobium
endo2 <- subset_taxa(ps_prunedSet, Genus=="Candidatus_Endomicrobium")
endo2

# Extract abundance matrix from the phyloseq object
OTU_endo2 = as(otu_table(endo2), "matrix")
# transpose if necessary
if(taxa_are_rows(endo2)){OTU_endo2 <- t(OTU_endo2)}
# Coerce to data.frame
OTU_endo2_df = as.data.frame(OTU_endo2)
write.table(OTU_endo2_df, file = "", sep = " ")

# Write taxonomy file
tax_prundedset <- tax_table(ps_prunedSet)

write.csv(tax_prundedset, file = "", sep = " " )

### Rarefaction curves #####
ggrare(ps_prunedSet, step = 20, se = TRUE, label = "cell_id")+
  facet_wrap(~Host_genus)
ggsave("rare_graph.png", path = "" )

#Remove Endomicrobium Sequences
# Define Genera to filter
filterEndos = c("Candidatus_Endomicrobium")

# Filter entries with Genera list from above
ps_prunedSet_noEndos = subset_taxa(ps_prunedSet, !Genus %in% filterEndos)
ps_prunedSet_noEndos

ggrare(ps_prunedSet_noEndos, step = 10, se = TRUE, label = "cell_id")+
  facet_wrap(~Host_genus)
ggsave("rare_noEndos_graph.png", path = "" )

#Revomve Endo only samples

ps_prunedSet_noEndos_b = subset_samples(ps_prunedSet_noEndos, cell_id != "BP")
ps_prunedSet_noEndos_b

ggrare(ps_prunedSet_noEndos_b, step = 10, se = TRUE, label = "cell_id")+
  facet_wrap(~Host_genus)
ggsave("rare_noEndosB_graph.png", path = "" )
